# *Pseudomonas aeruginosa* deploys competitor-specific antagonistic strategies

**DOI:** 10.64898/2026.04.14.718318

**Authors:** Ziyang Chen, Ned S. Wingreen, Zemer Gitai

**Affiliations:** Department of Molecular Biology, Princeton University, Princeton, New Jersey, USA; Lewis-Sigler Institute for Integrative Genomics, Princeton University, Princeton, New Jersey, USA

## Abstract

Microbial competition shapes polymicrobial communities, yet it remains unclear whether bacteria deploy fixed or specific antagonistic strategies against different rivals. Here we show that *Pseudomonas aeruginosa* deploys distinct strategies to outcompete two clinically relevant species, *Burkholderia cenocepacia* and *Staphylococcus aureus*. Under matched conditions, competition with *B. cenocepacia* is mediated by contact-dependent Type VI secretion, whereas competition with *S. aureus* follows a staged diffusible program of alkyl quinolone-mediated growth inhibition followed by LasA-dependent lysis. Transcriptomic analysis supports a “Swiss Army knife” model of antagonism in which different competitive context triggers distinct subsets of *P. aeruginosa*’s arsenal. Minimal dynamical models verify that contact-dependent killing is effective against slower-growing competitors, whereas a staged strategy of growth inhibition followed by killing via diffusible factors is preferable against faster-growing rivals. Together, these results show that the competitive success of *P. aeruginosa* depends on competitor-specific antagonistic strategies.

## 2 Introduction

Microbial communities in natural and clinical environments are inherently multi-species, with organisms constantly interacting through competition and cooperation [1, 2]. Of the two, competition is the dominant force shaping the structure, stability, and evolutionary trajectories of communities [3, 4]. Competition can occur indirectly through the consumption of shared resources, or directly via active antagonistic interactions. Although numerous antagonistic mechanisms have been identified across bacterial species, our mechanistic understanding of how inter-species competition unfolds in time and space, and whether bacteria deploy different antagonistic strategies in response to specific competitors, remains limited [5, 6].

*Pseudomonas aeruginosa* is a paradigmatic example of a bacterial species that is highly competitive in polymicrobial settings. As an opportunistic pathogen it frequently emerges as a dominant member of diverse communities, including those associated with chronic infections [7–9]. *P. aeruginosa*’s competitive success has been attributed to its unusually large arsenal of antagonistic mechanisms, encompassing diffusible small molecules, secreted enzymes, and contact-dependent killing systems [10, 11]. Extensive work has characterized individual components of this arsenal, revealing their molecular mechanisms and regulatory pathways. However, the presence of many antagonistic tools raises a fundamental question: Are these mechanisms differentially deployed during competition with different microbial neighbors? In other words, does *P. aeruginosa* employ a fixed set of antagonistic responses, or does it dynamically select, prioritize, and stage distinct strategies based on the identity and properties of competing species? This key question is unresolved because most prior studies focused on individual antagonistic mechanisms in isolation or examined interactions with a single competitor [5, 12–14], making it difficult to conclude whether *P. aeruginosa* employs competitor-dependent strategies under otherwise comparable conditions.

Here, we systematically examine how *P. aeruginosa* competes against two ecologically and physiologically distinct bacterial species: *Staphylococcus aureus*, a fast-growing Gram-positive bacterium [15], and *Burkholderia cenocepacia*, a metabolically versatile Gram-negative bacterium with substantial intrinsic antibiotic resistance [16]. Both species frequently co-occur with *P. aeruginosa* in polymicrobial infections [17–22]. Under matched experimental conditions, we found that *P. aeruginosa* deploys distinct antagonistic programs against different competitors. Specifically, we integrated spatial competition assays on agar, liquid coculture measurements, microscopy imaging, transcriptomic profiling, and targeted genetic perturbations to identify a Type VI secretion system that is specifically induced in the presence of *B. cenocepacia* to induce contact-dependent killing and alkyl quinolone and elastase diffusible factors that act sequentially to first arrest and then kill *S. aureus* at a distance. We further found that growth-rate asymmetry and resource allocation shape the effectiveness of these alternative strategies by developing minimal dynamical models that generalize our specific findings. Together, this combined experimental and theoretical approach reveals that *P. aeruginosa* uses its arsenal strategically, providing insight into how a single microbial species can flexibly dominate diverse opponents.

## 3 Results

### 3.1 Distinct interaction dynamics for *P. aeruginosa* competing with *S. aureus* versus *B. cenocepacia*

To frame how *P. aeruginosa* responds to different competitors, we conceptualized two alternative models: a “kitchen sink” model, in which *P. aeruginosa* deploys the same, full antagonistic repertoire against all competitors, and a “Swiss Army knife” model, in which *P. aeruginosa* selectively engages different strategies depending on the competing species (Fig. 1A). To distinguish between these possibilities, we compared *P. aeruginosa* competitions against *S. aureus* and *B. cenocepacia* (referred to as *Pa*:*Sa* and *Pa*:*Bc* respectively) under matched experimental conditions, focusing on how antagonism unfolds across space and time.

**Figure 1.**
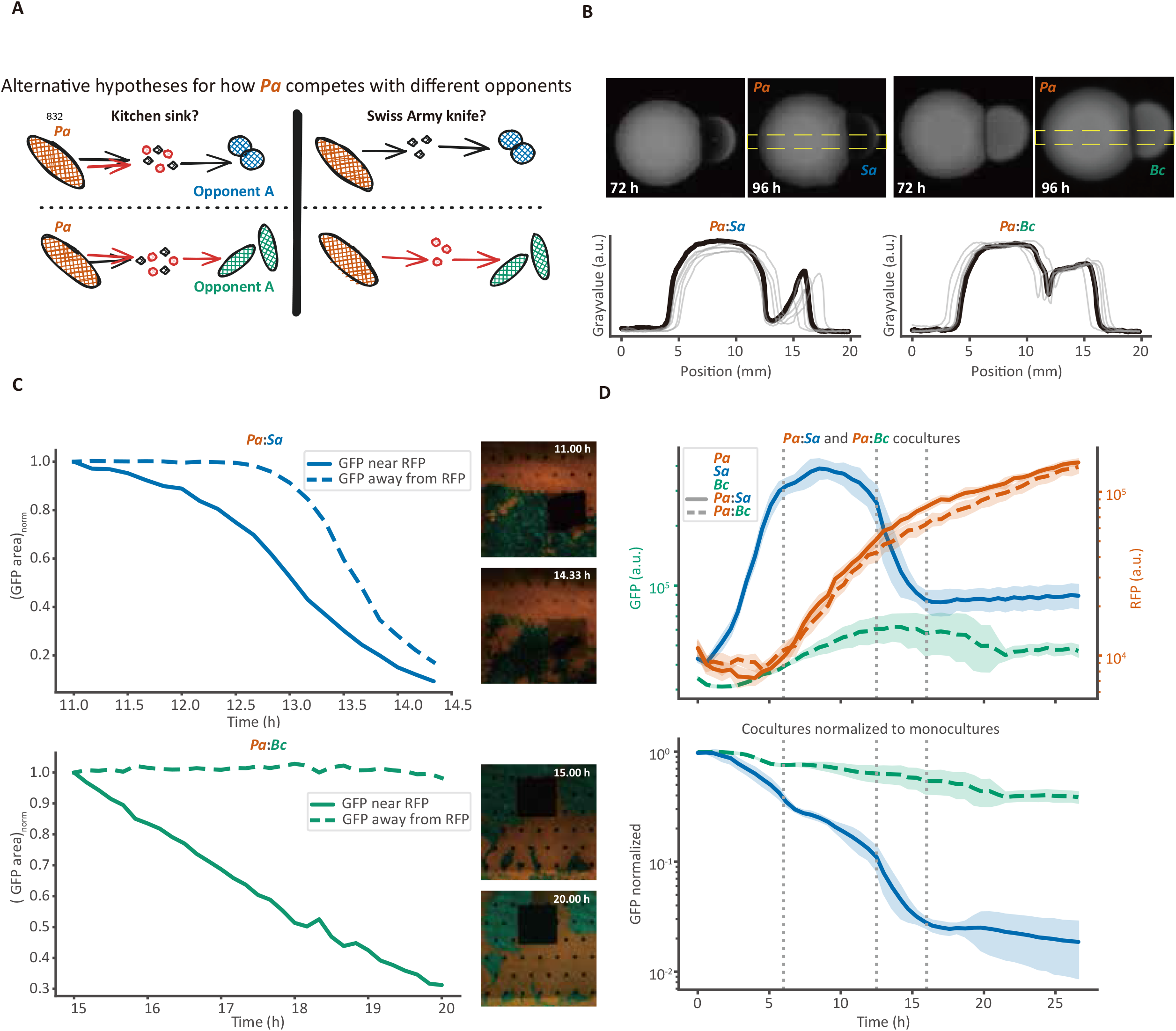
Distinct interaction dynamics of *P. aeruginosa* competing with *S. aureus* versus *B. cenocepacia*. (A) Two possible scenarios for how *P. aeruginosa* (*Pa*) competes against different opponents. (B) Spatial competition on solid media. Monocultures of *Pa*, together with *S. aureus* (*Sa*) or *B. cenocepacia* (*Bc*) were inoculated adjacently on LB agar plates and incubated at 37 °C for 96 h. Representative images (top) and corresponding grayscale intensity profiles across colonies (bottom; *n* = 6) are shown; the dashed yellow box indicates the region used for quantification. (C) Spatial dynamics in microfluidic coculture. *P. aeruginosa* (labeled with RFP) and its competitor bacteria (labeled with GFP) were grown together, and their interactions were observed over time by imaging. GFP area was defined as the number of pixels exceeding a threshold and normalized to the initial time point. (D) Bulk coculture dynamics in liquid medium. Bacteria were grown in 96-well plates, and fluorescence was monitored over time using a plate reader (*n* = 9). The upper panel shows fluorescence intensity dynamics of *P. aeruginosa* with *S. aureus* or with *B. cenocepacia* in coculture. The lower panel shows normalized fluorescence dynamics of *S. aureus* and *B. cenocepacia* in cocultures relative to their respective monocultures.

On agar surfaces, the two competitions exhibited markedly different spatial patterns of suppression. When co-inoculated adjacent to each other, *P. aeruginosa* inhibited *S. aureus* over a broad region extending well beyond the inter-colony interface, with *S. aureus* growth suppressed at substantial distances from *P. aeruginosa* (Fig. 1B). In contrast, during competition with *B. cenocepacia*, killing by *P. aeruginosa* was confined to a narrow zone at the inter-colony interface: *B. cenocepacia* remained largely unaffected in regions away from *P. aeruginosa* (Fig. 1B). Thus, even in a simple spatial assay, the pattern of suppression differed qualitatively depending on the opponent.

To learn how these spatial differences develop dynamically, we performed time-resolved single-cell-resolution imaging in microfluidic chambers, tracking fluorescently labeled populations (RFP labeled *P. aeruginosa*, GFP labeled *S. aureus* or *B. cenocepacia*) over time in cocultures. In *Pa*:*Sa* cocultures, *S. aureus* populations declined over time both near to and far from *P. aeruginosa* populations, indicating that killing was not confined to regions of direct contact (Fig. 1C, Extended Data Movie 1). In contrast, in *Pa*:*Bc* cocultures, *B. cenocepacia* loss occurred exclusively in regions immediately adjacent to *P. aeruginosa* populations, while distal populations were unaffected (Fig. 1C, Extended Data Movie 2). Together, these observations revealed that the two cocultures differ not only in spatial structure but also in the dynamics of killing: in *Pa*:*Sa* competition, *S. aureus* was eliminated across a broad spatial range with an abrupt collapse, whereas in *Pa*:*Bc* competition, *B. cenocepacia* killing was confined to the interface and proceeded in a slower, more continuous manner.

We next asked whether these distinct temporal interaction patterns are reflected in population-level dynamics in liquid. Using plate-reader-based fluorescence measurements, we monitored coculture trajectories over extended time courses. In *Pa*:*Sa* cocultures, the *S. aureus* GFP signal exhibited a characteristic three-phase behavior, with an initial growth phase, followed by a plateau, and then a rapid decline (Fig. 1D). In contrast, *Pa*:*Bc* cocultures displayed qualitatively different fluorescence dynamics, characterized by slower initial growth and persistent suppression relative to monocultures, but notably lacking the abrupt decline observed in *Pa*:*Sa* cocultures (Fig. 1D).

Together, these observations demonstrate that the antagonistic dynamics between *P. aeruginosa* and its competitors differ markedly in both spatial reach and temporal dynamics. These differences, observed across assays and scales, raise the question of whether they arise from competitor-specific antagonistic strategies deployed by *P. aeruginosa* (“Swiss Army knife”), or instead reflect differential susceptibilities of *S. aureus* and *B. cenocepacia* to “kitchen sink” antagonism. Therefore, we next examined whether these distinct interaction dynamics are accompanied by divergent transcriptional responses of *P. aeruginosa*.

### 3.2 Distinct transcriptomic responses of *P. aeruginosa* in coculture with *S. aureus* versus *B. cenocepacia*

The qualitatively distinct spatial and temporal interactions observed in Fig. 1 raise the possibility that *P. aeruginosa* engages different transcriptional programs depending on the identity of its competitor. To test this directly, we compared the transcriptomic responses of *P. aeruginosa* during coculture with *S. aureus* (*Pa*:*Sa*) or *B. cenocepacia* (*Pa*:*Bc*) under matched conditions, quantifying differential gene expression relative to *P. aeruginosa* in monoculture.

To directly assess the similarity or divergence of *P. aeruginosa* gene expression between the two coculture conditions, we plotted the log_2_ fold change (log_2_FC) of each *P. aeruginosa* gene in *Pa*:*Sa* against its log_2_FC in *Pa*:*Bc* (Fig. 2A). Genes with small expression changes under both conditions (|log_2_FC| < 1) clustered near the origin and were classified as not differentially expressed. Among genes that were differentially expressed in at least one coculture condition, only a minority exhibited concordant regulation: 7.3% were upregulated in both *Pa*:*Sa* and *Pa*:*Bc*, and 4.1% were downregulated in both conditions. In contrast, the vast majority (88.6%) displayed condition-specific regulation, showing substantial expression changes in one coculture but not the other, or changing in opposite directions. Thus, most of the transcriptional responses elicited during coculture were specific to a single competitor.

**Figure 2.**
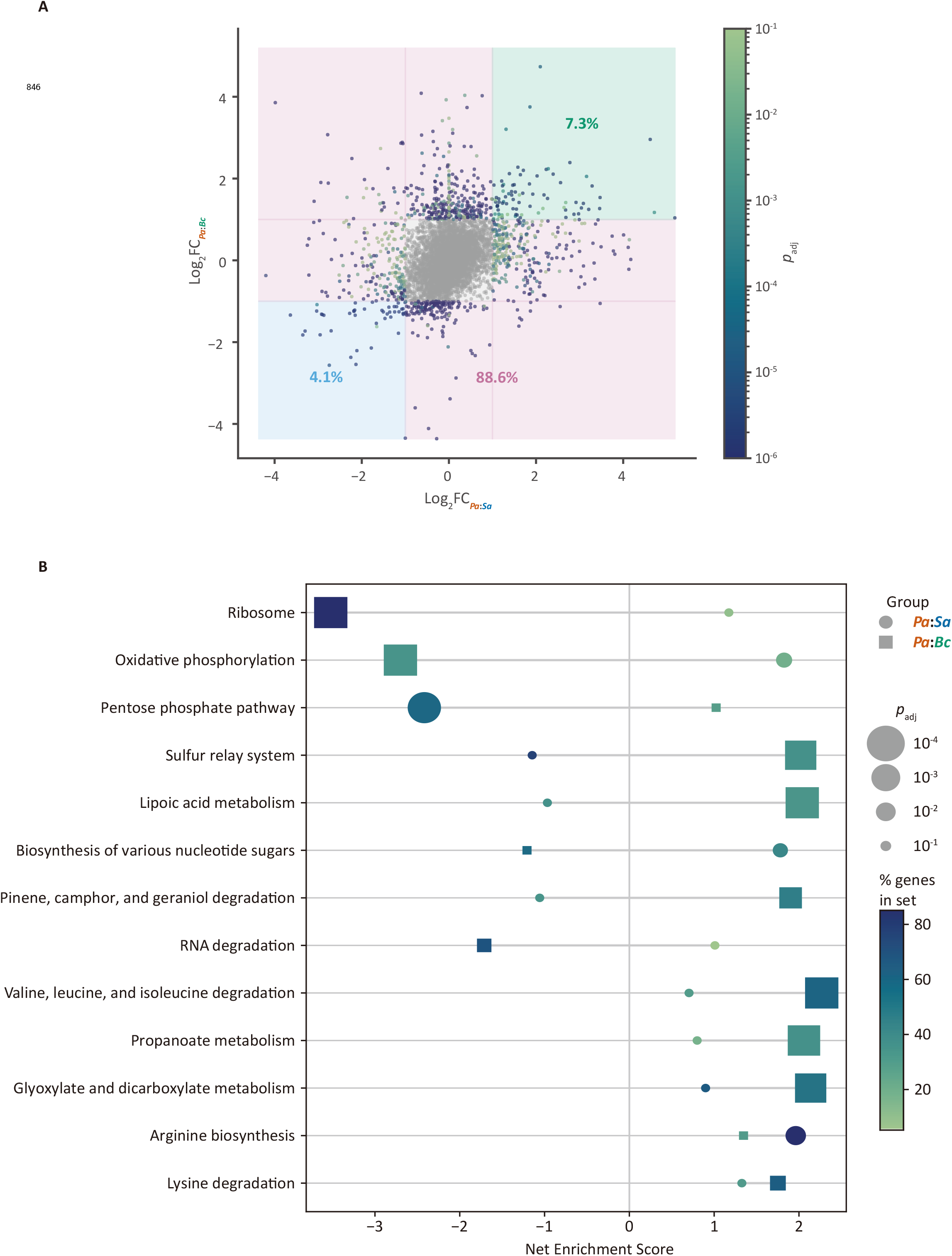
Distinct transcriptomic responses of *P. aeruginosa* in coculture with *S. aureus* versus *B. cenocepacia*. (A) Differential gene expression of *P. aeruginosa* during cocultures with *S. aureus* versus *B. cenocepacia*, with log_2_ fold changes calculated relative to *P. aeruginosa* monocultures (*n* = 3). Genes are colored according to adjusted *p*-values (FDR-corrected). Among genes differentially expressed in at least one condition (*Pa*:*Sa* or *Pa*:*Bc*), 7.3% (green region) were concordantly upregulated and 4.1% (blue region) concordantly downregulated in both cocultures, whereas 88.6% (pink regions) exhibited condition-specific regulation. (B) Gene Set Enrichment Analysis (GSEA) of *P. aeruginosa* transcriptomes in the two coculture conditions. Gene sets were defined using the Kyoto Encyclopedia of Genes and Genomes (KEGG) database, and enrichment was assessed using the GSEA pre-ranked method. The percentage of genes in set indicates how much of each pathway is represented by the genes driving the enrichment. The FDR-adjusted *p*-value (*p*_adj_) reflects the likelihood that an observed pathway enrichment occurs by chance.

Next, we asked how this transcriptional divergence is reflected at the level of functional gene sets. We performed Gene Set Enrichment Analysis (GSEA [23]) using KEGG-defined (Kyoto Encyclopedia of Genes and Genomes [24]) gene sets and compared normalized enrichment scores (NES) between the two coculture conditions (Fig. 2B). Many pathways exhibited markedly different enrichment patterns between the two cocultures, with some gene sets enriched in one condition but depleted or not enriched in the other. Notably, this divergence was observed across a broad range of functional categories, including core cellular processes and central metabolic pathways. These pathway-level differences indicate that *P. aeruginosa* undergoes extensive functional reprogramming that specifically depends on the competitor it encounters.

Together, this gene- and pathway-level analysis demonstrates that *P. aeruginosa* mounts largely distinct transcriptomic responses during coculture with *S. aureus* versus *B. cenocepacia*. This extensive divergence supports a competitor-dependent, “Swiss Army knife” model of antagonism and suggests that the different interaction dynamics observed in Fig. 1 arise from fundamentally different underlying regulatory states. We were thus motivated to undertake a mechanistic examination of the specific strategies deployed against each competitor.

### 3.3 *P. aeruginosa* outcompetes *B. cenocepacia* via Type VI secretion-mediated contact-dependent killing

The spatially restricted killing of *B. cenocepacia* observed in agar assay and microfluidic time-lapse imaging (Fig. 1B,C) suggests that *P. aeruginosa*-mediated antagonism against *B. cenocepacia* requires direct cell-cell contact. Building on this observation, and motivated by the competitor-specific transcriptional programs identified in Fig. 2, we sought to identify the mechanism responsible for *P. aeruginosa*-mediated suppression of *B. cenocepacia*.

To distinguish between contact-dependent and diffusible killing mechanisms, we first tested whether *B. cenocepacia* growth could be inhibited by spent coculture medium in the absence of *P. aeruginosa*. Whereas *S. aureus* failed to grow in spent medium derived from *Pa*:*Sa* cocultures, *B. cenocepacia* grew robustly in *Pa*:*Bc* coculture spent medium (Fig. 3A). These results indicate that, unlike suppression of *S. aureus*, antagonism against *B. cenocepacia* is not mediated by stable diffusible factors and instead requires the physical presence of *P. aeruginosa*.

**Figure 3.**
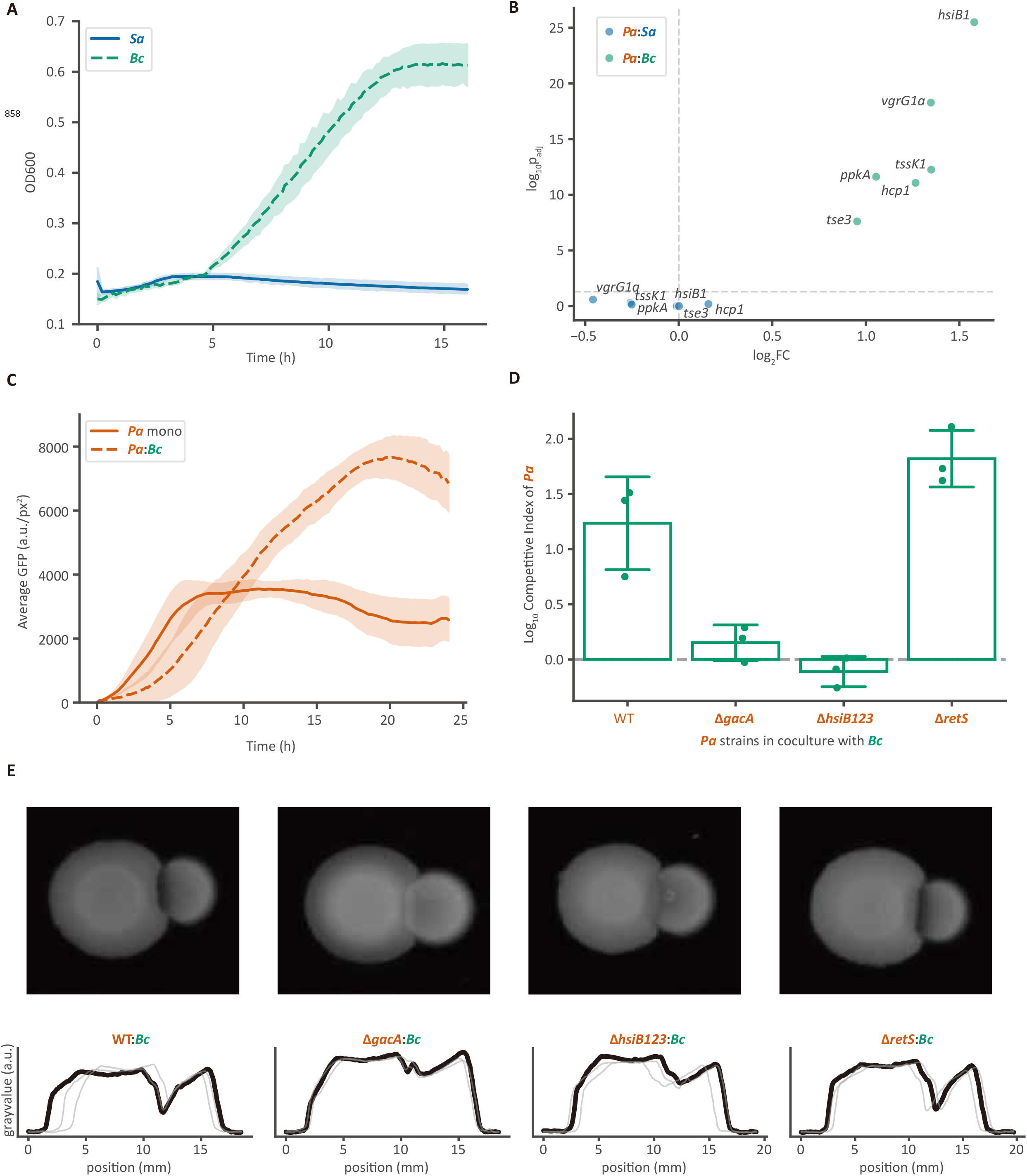
*P. aeruginosa* outcompetes *B. cenocepacia* via Type VI-mediated killing. (A) Growth of *S. aureus* and *B. cenocepacia* in respective coculture spent media in the absence of *P. aeruginosa* (*n* = 3). Coculture spent media were diluted 1:3 with fresh LB medium (spent medium:LB). (B) Differential expression of representative Type VI secretion system (T6SS) genes in *P. aeruginosa* during coculture with *S. aureus* (*Pa*:*Sa*) or *B. cenocepacia* (*Pa*:*Bc*) after 2.5 h. log_2_ fold changes were calculated relative to *P. aeruginosa* monocultures (*n* = 3). (C) Average fluorescence dynamics of an *P. aeruginosa hsiB1* translational reporter strain (Δ*retS* background) grown in the presence or absence of *B. cenocepacia*. Fluorescence was monitored by time-lapse imaging (*n* = 5). Average GFP was calculated by normalizing total GFP signal by area (1 px=65 nm). (D) Competitive indices (see Methods) of *P. aeruginosa* strains after 6 h of coculture with GFP-labeled *B. cenocepacia*. Cocultures were initiated by mixing *P. aeruginosa* with *B. cenocepacia* and spotting onto LB agar plates (*n* = 3). Competitive indices were calculated after 6 h of growth at 37 °C. (E) Agar plate competition assays between *B. cenocepacia* and *P. aeruginosa* using *P. aeruginosa* deletion mutants. Representative images (top) and corresponding grayscale intensity profiles across colonies (bottom; *n* = 3) are shown.

Relatively few contact-dependent antagonistic mechanisms have been described for *P. aeruginosa* [25–27], among which the Type VI Secretion System (T6SS) is a well-established example [28, 29]. Consistent with a contact-dependent mode of killing, the transcriptomic analysis revealed strong and specific upregulation of T6SS genes in *P. aeruginosa* during coculture with *B. cenocepacia*, but not during coculture with *S. aureus* (Fig. 3B). To directly assess the time course of T6SS activity, we used a translational reporter for the T6SS contractile sheath component *hsiB1* in a Δ*retS* background. (This background places cells in a T6SS-permissive regulatory state and improves the dynamic range of reporter detection by relieving RetS-mediated repression of the Gac/Rsm pathway [30, 31].) In this sensitized background, reporter fluorescence increased markedly in the presence of *B. cenocepacia*, but not in monoculture, confirming that the presence of *B. cenocepacia* stimulates T6SS expression in *P. aeruginosa* (Fig. 3C).

We next tested whether T6SS activity is required for effective killing of *B. cenocepacia*. To do so, we performed well-mixed coculture competition assays and measured the competitive index (see Methods) of wild-type *P. aeruginosa* and a set of deletion mutants. Disrupting key regulatory or structural components of the T6SS significantly weakened *P. aeruginosa*’s ability to suppress *B. cenocepacia*. Specifically, mutants lacking *gacA*, a positive regulator of T6SS, or all three *hsiB* loci, which encode contractile sheath components, exhibited substantially lower competitive indices than wild-type, whereas deletion of *retS*, a negative regulator of T6SS, increased the competitive index (Fig. 3D). Notably, these mutants remained as competitive as wildtype in coculture with *S. aureus*, indicating that the impaired suppression of *B. cenocepacia* arises from a specific defect in *Pa*:*Bc* antagonism rather than from a general loss of competitive fitness (Extended Data Fig. 1A).

These genetic requirements were also mirrored in spatial competition assays. Wild-type *P. aeruginosa* and the Δ*retS* mutant produced a zone cleared of *B. cenocepacia* at the colony interface, whereas Δ*gacA* and Δ*hsiB123* mutants failed to generate a detectable killing zone (Fig. 3E). Quantitative intensity profiles across the colony boundary confirmed the loss of interface-restricted killing in T6SS-deficient strains. Importantly, the same mutants did not exhibit analogous defects in agar-based competition with *S. aureus*, further demonstrating that T6SS-dependent contact killing is selectively deployed during competition with *B. cenocepacia* (Extended Data Fig. 1C).

Together, these findings provide a mechanistic explanation for the spatially restricted suppression of *B. cenocepacia* observed in *Pa*:*Bc* cocultures in Fig. 1. Specifically, they demonstrate that *P. aeruginosa* suppresses *B. cenocepacia* through contact-dependent, T6SS-mediated killing, explaining why antagonism is confined to regions of direct physical proximity. This mode of competition contrast sharply with the long-range and temporally structured suppression of *S. aureus* in *Pa*:*Sa* cocultures, which we examine next.

### 3.4 *P. aeruginosa* employs a stepwise brake-then-break diffusible strategy to outcompete *S. aureus*

Unlike the contact-dependent killing of *B. cenocepacia*, competition between *P. aeruginosa* and *S. aureus* (Fig. 1) exhibited long-range inhibition and a distinct temporal structure, including an extended plateau followed by a rapid decline in *S. aureus* abundance. Consistent with these distinct features, genome-wide expression profiling (Fig. 2) indicated that *P. aeruginosa* enters a specialized regulatory state during coculture with *S. aureus*. Together, these observations suggest that *P. aeruginosa* suppresses *S. aureus* through a temporally structured strategy that is mechanistically distinct from contact-dependent killing. We therefore sought to identify the functional basis of *P. aeruginosa*-mediated antagonism against *S. aureus*.

To investigate *P. aeruginosa*’s antagonistic program during *Pa*:*Sa* competition, we focused on functions that could account for the three-phase dynamics observed in Fig. 1D. The plateau phase suggests growth suppression, whereas the subsequent rapid decline implies a more potent killing mechanism. Consistent with this logic, *P. aeruginosa* is known to secrete several respiration-inhibiting small molecules (including those regulated by the PQS quorum-sensing system [32]), and also to secrete a peptidoglycan hydrolase LasA, which has staphylolytic activity [33]. We therefore examined the expression dynamics of *pqsA*, a key gene in PQS biosynthesis, and of *lasA* during coculture with *S. aureus*. Transcriptomic analysis revealed that *pqsA* was specifically upregulated at early time points during *Pa*:*Sa* coculture, whereas *lasA* induction was delayed and occurred predominantly at later stages (Fig. 4A). Notably, neither gene showed comparable induction during coculture with *B. cenocepacia*.

**Figure 4.**
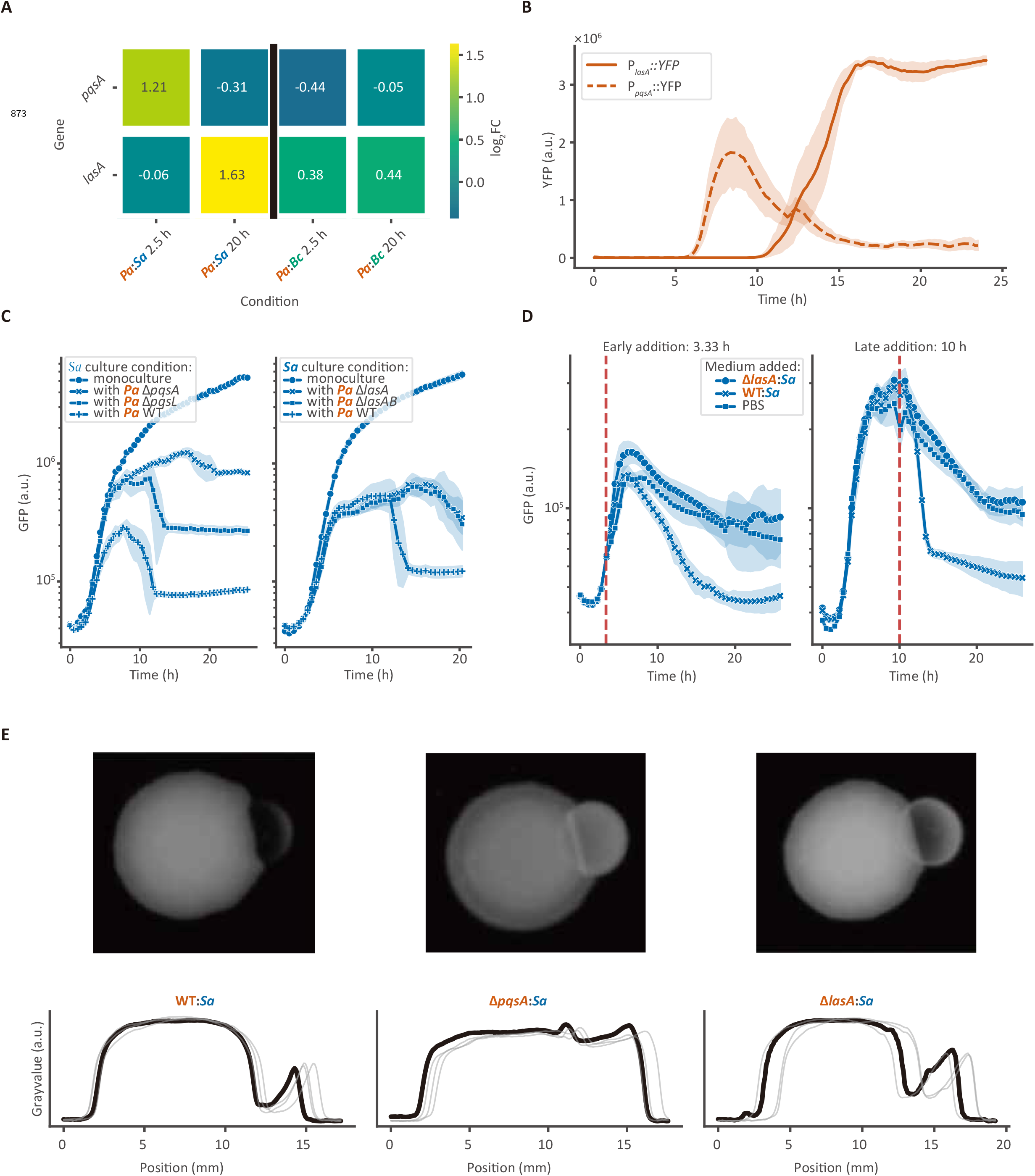
*P. aeruginosa* employs a stepwise, brake-then-break strategy to outcompete *S. aureus*. (A) Differential expression of *pqsA* and *lasA* in *P. aeruginosa* during coculture with *S. aureus* (*Pa*:*Sa*) or *B. cenocepacia* (*Pa*:*Bc*) at early (2.5 h) and late (20 h) time points. log_2_ fold changes were calculated relative to time-matched *P. aeruginosa* monocultures (*n* = 3). (B) Fluorescence dynamics of *pqsA* and *lasA* transcriptional reporter strains grown in *Pa*:*Sa* coculture spent medium. Coculture spent medium was diluted 1:3 with fresh LB medium (spent medium:LB). YFP fluorescence was monitored by timelapse imaging (*n* = 3). (C) Fluorescence dynamics of GFP-labeled *S. aureus* in coculture with *P. aeruginosa* deletion mutants (*n* = 9). Left, coculture with Δ*pqsA*, Δ*pqsL*, and wild-type (WT) *P. aeruginosa*. Right, coculture with Δ*lasA*, Δ*lasAB*, and WT *P. aeruginosa*. (D) Fluorescence dynamics of GFP-labeled *S. aureus* in coculture with Δ*lasA P. aeruginosa* supplemented with spent media (*n* = 9). Cocultures were supplemented with Δ*lasA*:*Sa* spent medium, WT:*Sa* spent medium, or PBS. Spent medium added either 3.33 h (red dashed vertical line) after coculture initiation (left) or 10 h after coculture initiation (right). (E) Agar plate competition assays between *P. aeruginosa* and *S. aureus* using *P. aeruginosa* deletion mutants. Representative images (top) and corresponding grayscale intensity profiles across colonies (bottom; *n* = 4) are shown.

Next, we used transcriptional reporter strains to directly monitor the dynamics of *pqsA* and *lasA* expression in response to *S. aureus*-conditioned environments. Previous studies have shown that diffusible molecules produced by *S. aureus* can modulate *P. aeruginosa* virulence gene expression [34, 35], motivating the use of *Pa*:*Sa* coculture spent medium as a proxy for *S. aureus*-associated cues. We found that exposure to spent medium induced early expression of the *pqsA* reporter, whereas expression of the *lasA* reporter increased only at later times (Fig. 4B). These results demonstrate that in response to *S. aureus, P. aeruginosa* activates PQS-associated responses prior to inducing LasA expression, consistent with a temporally ordered antagonism program.

To test whether these temporally staged programs correspond to the observed different phases of *S. aureus* suppression, we examined coculture dynamics using *P. aeruginosa* deletion mutants. In liquid coculture assays, deletion of *pqsA* or *pqsL* (which encodes a critical enzyme in the biosynthesis of respiration inhibitor 2-heptyl-4-hydroxyquinoline N-oxide, HQNO, [36]) abolished the early inhibitory phase observed in wild-type *Pa*:*Sa* cocultures, resulting in sustained *S. aureus* growth during the initial period (Fig. 4C, left). In contrast, deletion of *lasA* or *lasAB* specifically impaired the later killing phase without affecting early inhibition, leading to persistence of *S. aureus* at late time points (Fig. 4C, right). Similar defects in competitive index were also observed in well-mixed coculture competition assays between *P. aeruginosa* and *S. aureus* (Extended Data Fig. 1A). However, these mutants did not exhibit comparable defects in *Pa*:*Bc* cocultures (Extended Data Fig. 1B), indicating that PQS- and LasA-dependent antagonism is specific to the *Pa*:*Sa* interaction.

We next asked whether diffusible factors that accumulate in *Pa*:*Sa* coculture are sufficient to trigger killing of *S. aureus* independently of continued coculture. To address this, we supplemented Δ*lasA Pa*:*Sa* cocultures with spent medium collected from late-stage wild-type cocultures, in which inhibitory factors and LasA have accumulated to high levels, as indicated by growth of *S. aureus* and a staphylolysis assay (Extended Data Fig. 4A). Addition of this late-stage coculture spent medium early after coculture initiation abrogated the plateau phase and restored *S. aureus* killing, whereas addition at later time points triggered the immediate onset of killing (Fig. 4D). In contrast, PBS or spent medium derived from Δ*lasA* cocultures failed to restore killing. These results indicate that the accumulated diffusible components present in *Pa*:*Sa* coculture supernatants are sufficient to drive killing and that the timing of their availability governs the overall dynamics of *S. aureus* suppression.

Finally, we assessed whether these genetic dependencies are reflected in spatial competition assays. In agar-based competitions, wild-type *P. aeruginosa* produced a pronounced zone of *S. aureus* clearance that extended beyond the colony interface (Fig. 4E). In contrast, Δ*pqsA* and Δ*lasA* mutants exhibited marked defects in *S. aureus* killing, with reduced or absent clearance zones (Fig. 4E). These phenotypes were specific to *Pa*:*Sa* competition and were not observed in analogous assays with *B. cenocepacia* (Extended Data Fig. 1D), further underscoring the interaction-specific nature of this antagonistic strategy.

Together, these findings demonstrate that *P. aeruginosa* suppresses *S. aureus* through a stepwise antagonistic strategy, consisting of an early inhibitory phase followed by a killing phase. This temporally ordered mechanism provides a mechanistic “brake then break” explanation for the long-range multi-phase interaction dynamics observed in which *P. aeruginosa* first arrests and later lyses *S. aureus*.

### 3.5 Growth-rate asymmetry and metabolic costs shape antagonistic strategies

Our experimental results revealed that *P. aeruginosa* deploys fundamentally different antagonistic strategies depending on competitor identity: contact-dependent T6SS activity dominates during competition with *B. cenocepacia*, whereas diffusible inhibition followed by lytic enzyme-mediated killing characterizes interactions with *S. aureus*. These two competitors also differ markedly in intrinsic growth rate, with *B. cenocepacia* growing at approximately the same rate as *P. aeruginosa*, but *S. aureus* growing substantially faster (Extended Data Fig. 2A). This raised the possibility that the observed divergence in antagonistic programs does not simply reflect arbitrary species-specific molecular recognition, but instead emerges from fundamental considerations dictated by growth-rate asymmetry and resource competition. To test this hypothesis, we constructed minimal dynamical models of contact-dependent and diffusible antagonism to determine how competitor growth rate and resource allocation shape the relative efficacies of distinct killing strategies.

We first asked under what conditions is contact-dependent antagonism alone sufficient to confer a competitive advantage? To isolate the essential dynamics of T6SS-mediated interactions, we implemented a minimal 2D model in which killing occurs exclusively upon direct cell–cell contact, while both populations – *P. aeruginosa* and competitor – compete for a shared limiting resource. Simulations revealed a sharp dependence on growth-rate asymmetry: when the competitor grows at a rate equal to or slower than *P. aeruginosa*, contact-dependent killing effectively suppresses the former’s expansion and enables *P. aeruginosa* to dominate. In contrast, when the competitor’s intrinsic growth rate exceeds that of *P. aeruginosa*, competitor population expansion outpaces its contact-mediated killing, preventing *P. aeruginosa* from dominating (Fig. 5A and Fig. 5B). At higher growth-rate ratios the variability of simulation results (due to random initial cell placement) increases, which suggests that contact-dependent killing is not only less effective on average against faster-growing competitors, but also less reliable. These results indicate that contact-dependent killing is intrinsically limited against faster-growing species, providing a mechanistic explanation for why T6SS activity is effective against the slower-growing *B. cenocepacia* but is not employed by *P. aeruginosa* when faced with the more rapidly expanding *S. aureus*.

**Figure 5.**
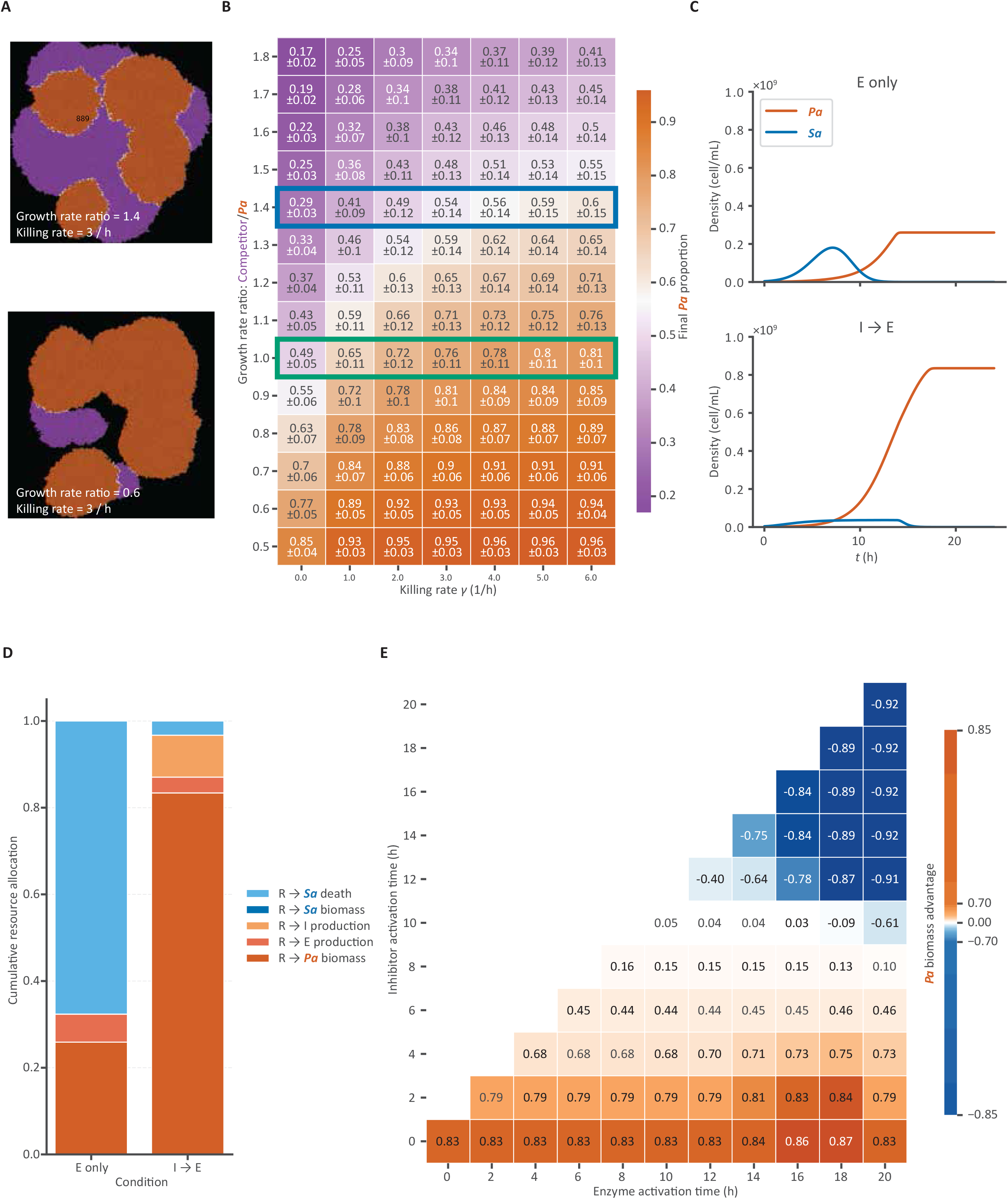
Growth-rate asymmetries and metabolic costs shape antagonistic strategies. (A) Representative snapshots of competition between *P. aeruginosa* (orange) and a competitor (purple) when 50% carrying capacity is reached in 2D simulations. (B) Proportion of *P. aeruginosa* at 50% carrying capacity across growth-rate ratios and killing rates (mean ± s.d., *n* = 9 independent simulations). The blue box indicates the growth rate ratio corresponding to *S. aureus* over *P. aeruginosa*, while the green box indicates the ratio corresponding to *B. cenocepacia* over *P. aeruginosa*. Simulations were initialized with 10 *P. aeruginosa* and 10 competitor cells randomly positioned in space. (C) Simulated bulk dynamics of *P. aeruginosa* and *S. aureus* under enzyme-only (E only) and sequential inhibitor-then-enzyme (I→E) strategies. In E only, enzyme activation occurs at 2 h with *A*_*E*_ = 0.3. In I→E, inhibitor and enzyme activation occur at 2 h and 10 h, with *A*_*I*_ = 0.2 and *A*_*E*_ = 0.1, respectively. (D) Cumulative resource allocation at 24 h for E only and I→E. Categories denote resources allocated to killed *S. aureus* biomass (*R* → *Sa* death), surviving *S. aureus* biomass (*R* → *Sa* biomass), inhibitor/enzyme production (*R* → *I/E* production), and *P. aeruginosa* biomass (*R* → *Pa* biomass). (E) Optimized final biomass advantage of *P. aeruginosa* as a function of inhibitor and enzyme activation times and allocation amplitudes (see Extended Data Fig. 3). Biomass advantage is defined as the final biomass difference between *P. aeruginosa* and *S. aureus*.

If the effectiveness of contact-dependent killing is intrinsically limited against a faster-growing opponent, then, faced with such a competitor, *P. aeruginosa* must rely on mechanisms that operate beyond immediate physical contact. Diffusible antagonistic molecules overcome this spatial constraint by acting over longer ranges. However, secreted factors must be expressed at high levels to maintain effective concentrations despite diffusion. Thus, employing diffusible growth inhibitors and/or lytic enzymes may well be more costly than production of contact-dependent-killing factors in terms of diverting metabolic flux away from biomass production. Indeed, production of diffusible factors could slow *P. aeruginosa* expansion at precisely the stage when resource capture is most critical, allowing the competitor to consume shared resources before killing becomes effective. We therefore next asked how investment in diffusible inhibition and enzymatic killing affects competition, and whether rational temporal structuring of these programs can overcome the disadvantage imposed by rapid competitor growth.

To determine how diffusible antagonism performs against a fast-growing competitor, we compared two allocation strategies in a bulk resource-competition model: investment in lytic-enzyme production alone (“E-only”) versus sequential deployment in which growth inhibition precedes enzymatic killing (“I→E”). In the E-only scenario, enzyme-mediated killing begins early, but *S. aureus* continues to proliferate during the initial phase, rapidly capturing shared resources while *P. aeruginosa* diverts substantial metabolic flux toward enzyme synthesis (Fig. 5C). As a result, a substantial flux of resources remains directed into *S. aureus* biomass, despite ongoing killing, thus limiting the final extent of *P. aeruginosa* expansion (Fig. 5D). In contrast (“I→E”), early production of a relatively cheap but effective diffusible inhibitor suppresses *S. aureus* growth before significant resource capture occurs. Subsequent activation of enzymatic killing then eliminates a growth-arrested small population of *S. aureus*, allowing *P. aeruginosa* to ultimately consume a larger fraction of the shared resource pool (Fig. 5D). These simulations indicate that against a fast-growing species, a “brake then break” strategy can outperform immediate investment in killing.

Because sequential deployment of inhibition followed by killing outperformed constitutive enzyme production, we next asked how competitive success depends quantitatively on the timing of these two programs. If early inhibition limits resource capture by *S. aureus*, whereas delayed killing minimizes premature metabolic expenditure, then the advantage of *P. aeruginosa* should exhibit a non-monotonic dependence on activation times. To test this, we systematically varied both the onset times (*t*_*X*_) and allocation amplitudes (*A*_*X*_) of inhibitor and lytic-enzyme production and measured the resulting final biomass difference between the two species. We found that maximal *P. aeruginosa* biomass advantage is achieved by a temporally ordered “brake then break” strategy, in which early growth inhibition of *S. aureus* is followed by delayed enzymatic killing (Fig. 5E and Extended Data Fig. 3). The observed optimal delay falls between too-early enzyme deployment, which continuously diverts resources away from *P. aeruginosa* even after *S. aureus* has been effectively eliminated, and too-late deployment which allows competitor biomass to persist because late-time resource depletion limits *P. aeruginosa*’s lytic-enzyme production (Extended Data Fig. 2C and Extended Data Fig. 2D). In summary, temporal separation of growth inhibition then killing is advantageous, highlighting the importance of dynamic allocation strategies to minimize metabolic costs.

Together, these simulations identify growth-rate asymmetry and resource allocation constraints as fundamental determinants of antagonistic strategy. Contact-dependent killing is effective only when the target expands at a rate comparable to or slower than *P. aeruginosa*, explaining its success against the slower-growing *B. cenocepacia*. In contrast, diffusible antagonism overcomes spatial limitations to suppress a faster-growing foe, but at a higher metabolic cost. A sequential combination of growth inhibition followed by enzymatic killing proves to be a cost-effective strategy. Thus, competitive success depends not only on which antagonistic factors are deployed, but on how they are temporally staged. These results provide a quantitative framework for understanding how *P. aeruginosa* tailors its competitive strategy in light of competitor characteristics.

## 4 Discussion

In this study, we demonstrate that *P. aeruginosa* engages a “Swiss Army knife” approach to competition, selectively activating specific antagonistic programs tailored to the identity of the opposing species, rather than deploying a fixed arsenal. Across matched spatial, temporal, genetic, and transcriptomic analyses, we found that competition with *B. cenocepacia* is dominated by contact-dependent, Type VI secretion-mediated killing, whereas competition with *S. aureus* is governed by a mechanistically distinct, “brake then break”, temporally ordered diffusible-factor strategy in which growth inhibition precedes killing. Our modeling rationalized these alternative programs: contact killing is most effective against slower-growing competitors, whereas the more costly sequential “brake then break” strategy is advantageous against faster-growing rivals. Together, these results argue that the antagonistic success of *P. aeruginosa* depends on its ability to deploy the right weapon with the right timing against the right competitor.

Our results suggest that the success of a generalist competitor such as *P. aeruginosa* lies not simply in possessing many antagonistic weapons, but in deploying them strategically according to ecological context. This shift in focus from arsenal composition to arsenal deployment has important implications for how microbial competition is understood in complex communities. In polymicrobial settings, competitive outcomes may not be determined solely by which antagonistic systems are encoded in a genome, but also by how those systems are dynamically regulated in response to the identity, physiology, and growth behavior of neighboring species. In this view, the ability of *P. aeruginosa* to thrive across diverse polymicrobial environments may stem from sensory, regulatory, and physiological versatility which together allow it to calibrate antagonistic strategy and timing to match the growth characteristics and vulnerabilities of its competitors. Our findings therefore support a broader view of microbial antagonism as a context-dependent and finely-tuned behavior.

Several limitations should be considered when interpreting our findings. Our experiments were performed under controlled *in vitro* conditions that enabled matched comparison of distinct antagonistic programs, but such conditions necessarily simplify the environments in which *P. aeruginosa* naturally competes. In host-associated settings, spatial confinement, host-derived nutrients, immune activity, and physical barriers are all likely to reshape both antagonistic deployment and competitive outcomes [37–39]. Also, long-term competitions are often stabilized by evolutionary adaptations on both sides that mitigate conflict, such as the selection of *S. aureus* small colony variants [40] or the downregulation of *P. aeruginosa* virulence factors [41]. In addition, we focused on pairwise interactions, whereas natural polymicrobial communities often involve multispecies consortia in which higher-order interactions may modify the dynamics observed here. Finally, our minimal theoretical models were designed to capture ecological principles rather than provide quantitative comparison with data. Accordingly, our overall findings should be viewed as establishing a conceptual exemplar of competitor-specific antagonism that captures key principles shaping these interactions, without attempting to fully reconstruct their native-environmental complexity.

Future work will be required to determine how *P. aeruginosa* senses competitor identity and translates that information into distinct antagonistic programs. The upstream cues responsible for this specificity may include physical contact, diffusible metabolites, nutrient limitations, or stress signals generated during competition. For *B. cenocepacia*, the requirement for contact suggests a mechanism of “tit-for-tat” retaliation [42, 43]. However, diffusible cues may also contribute. Possible candidates include interspecies cross-talk mediated by N-acylhomoserine lactones (AHLs) [44] and signaling through *Burkholderia* diffusible signal factor (BDSF) [45]. For *S. aureus*, peptidoglycan fragments released during cell wall turnover or lysis [35], specific *S. aureus* signaling molecules [46], and iron starvation signals induced by siderophore competition [47] are likely candidates. Defining the regulatory circuitry that links these inputs to T6SS activation versus diffusible inhibitor- and enzyme-based antagonism will be an important next step toward understanding how strategic deployment is implemented mechanistically. It will also be important to test whether the principles identified here continue to hold in more complex ecological settings, including multispecies communities, structured biofilms, and host-associated environments, where additional interactions may reshape both antagonistic decisions and competitive outcomes. Expanding this framework across a broader panel of competitors may further reveal whether tuning antagonism to ecological context is a widely shared feature of generalist pathogens like *P. aeruginosa*

## 5 Methods

### 5.1 Bacterial strains and growth conditions

All bacterial strains, plasmids, and primers used in this study are listed in Supplementary Table 1, Supplementary Table 2, and Supplementary Table 3. Unless otherwise indicated, overnight cultures were grown in LB medium at 37 °C with orbital shaking at 200 rpm. Spent medium was prepared by centrifugation of cultures at 7000 × *g* for 10 min, followed by collection of the supernatant using syringe filtration (0.2 µm pore diameter, Whatman, cat# WHA67802502).

**Table 1:** Strains used in this study.

| Background | Description | Resistance | Reference |
| --- | --- | --- | --- |
| *E. coli* HB101 | pRK2013 | KanR | [58] |
| *E. coli* Pir1 | pTNS2 | AmpR | [59] |
| *E. coli* Pir1 | pTNS3 | AmpR | [60] |
| *E. coli* S17-1 | pEXG2-$\Delta pqsL$ | GenR | this study |
| *E. coli* S17-1 | pEXG2-$\Delta pqsA$ | GenR | this study |
| *E. coli* S17-1 | pEXG2-$\Delta lasA$ | GenR | this study |
| *E. coli* S17-1 | pEXG2-$\Delta lasB$ | GenR | this study |
| *E. coli* S17-1 | pEXG2-$\Delta hsiB1$ | GenR | this study |
| *E. coli* S17-1 | pEXG2-$\Delta hsiB2$ | GenR | this study |
| *E. coli* S17-1 | pEXG2-$\Delta hsiB3$ | GenR | this study |
| *E. coli* S17-1 | pEXG2-$\Delta gacA$ | GenR | this study |
| *E. coli* SM10 | pEXG2-$\Delta tssM1\_PA14$ | GenR | [31] |
| *E. coli* SM10 | pEXG2-$tssB1-mNG\_PA14$ | GenR | [31] |
| *E. coli* SM10 | pEXG2-$\Delta retS\_PA14$ | GenR | [31] |
| *E. coli* DH5α | pEXG2 empty vector maintaining strain | GenR | [48] |
| *E. coli* S17-1 | Vector mobilization host |  | [61] |
| NCDC 7119 (E0-1 group) | *B. cenocepacia* wild type strain |  | ATCC 25608 |
| NCDC 7119 (E0-1 group) | *B. cenocepacia* attTn7::mini-Tn7-*eGFP* | KanR | this study |
| UCBPP-PA14 | *P. aeruginosa* wild type strain |  | [61] |
| UCBPP-PA14 | *P. aeruginosa* attTn7::Gm-*Pc-DsRed-Express2* | GenR | this study |
| UCBPP-PA14 | *P. aeruginosa lasA* transcriptional reporter strain | AmpR | this study |
| UCBPP-PA14 | *P. aeruginosa pqsA* transcriptional reporter strain | AmpR | this study |
| UCBPP-PA14 | *P. aeruginosa pqsA* deletion strain |  | this study |
| UCBPP-PA14 | *P. aeruginosa pqsL* deletion strain |  | this study |
| UCBPP-PA14 | *P. aeruginosa lasA* deletion strain |  | this study |
| UCBPP-PA14 | *P. aeruginosa lasA* and *lasB* deletion strain |  | this study |
| UCBPP-PA14 | *P. aeruginosa retS* deletion strain |  | this study |
| UCBPP-PA14 | *P. aeruginosa gacA* deletion strain |  | this study |
| UCBPP-PA14 | *P. aeruginosa hsiB123* deletion strain |  | this study |
| UCBPP-PA14 | *P. aeruginosa dretS hsiB1* translational reporter strain | GenR | this study |
| USA300 LAC | *S. aureus* wild type strain |  | [62] |
| USA300 LAC | *S. aureus* strain with pOS1-*PsarA-sodRBS-sGFP* | CmR | [62] |

**Table 2:** Plasmids used in this study.

| Name | Description | Resistance | Reference |
| --- | --- | --- | --- |
| pUC18R6KT-mini-Tn7T-*PS12-eGFP*-KAN | Integration vector for miniTn7 element containing *eGFP* and kanamycin resistance cassette | KanR | [63] |
| pTNS3 | Tn7 transposase expression | AmpR | [60] |
| pRK2013 | Helper plasmid for mobilization | KanR | [58] |
| pUC18T-mini-Tn7T-Gm-*Pc-DsRed-Express2* | mini-Tn7 vector for insertion of *DsRed-Express2* at bacterial attTn7 site | GenR, AmpR | [64] |
| pTNS2 | Tn7 transposase expression | AmpR | [59] |
| pEXG2 | Suicide vector for allele exchange | GenR | [48] |
| pUCP18-*P<sub>PaQa</sub>::mVenus-P<sub>rpoD</sub>::mKate* | *P. aeruginosa* transcriptional reporter template | AmpR | [65] |
| pUCP18-*P<sub>lasA</sub>::mVenus-P<sub>rpoD</sub>::mKate* | *lasA* transcriptional reporter | AmpR | this study |
| pUCP18-*P<sub>pqsA</sub>::mVenus-P<sub>rpoD</sub>::mKate* | *pqsA* transcriptional reporter | AmpR | this study |
| pEXG2-Δ *pqsL* | *pqsL* deletion plasmid | GenR | this study |
| pEXG2-Δ *lasA* | *lasA* deletion plasmid | GenR | this study |
| pEXG2-Δ *lasB* | *lasB* deletion plasmid | GenR | this study |
| pEXG2-Δ *tssM1_PA14* | *icmF1* deletion plasmid | GenR | [31] |
| pEXG2-*tssB1-mNG_PA14* | *hsiB1* translational reporter plasmid | GenR | [31] |
| pEXG2-Δ *retS_PA14* | *retS* deletion plasmid | GenR | [31] |
| pEXG2-Δ *gacA* | *gacA* deletion plasmid | GenR | this study |
| pEXG2-Δ *hsiB1* | *hsiB1* deletion plasmid | GenR | this study |
| pEXG2-Δ *hsiB2* | *hsiB2* deletion plasmid | GenR | this study |
| pEXG2-Δ *hsiB3* | *hsiB3* deletion plasmid | GenR | this study |

**Table 3:**
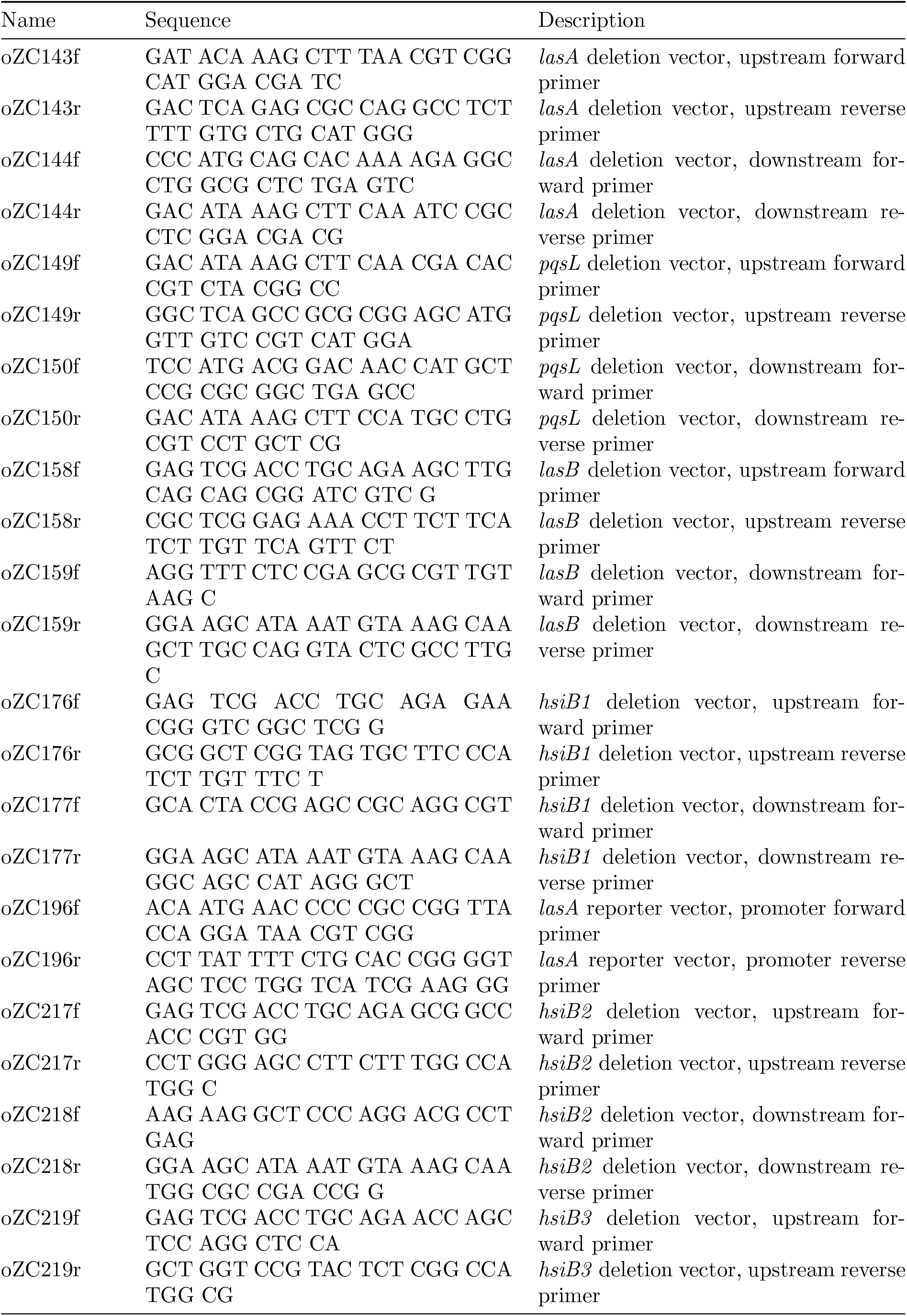
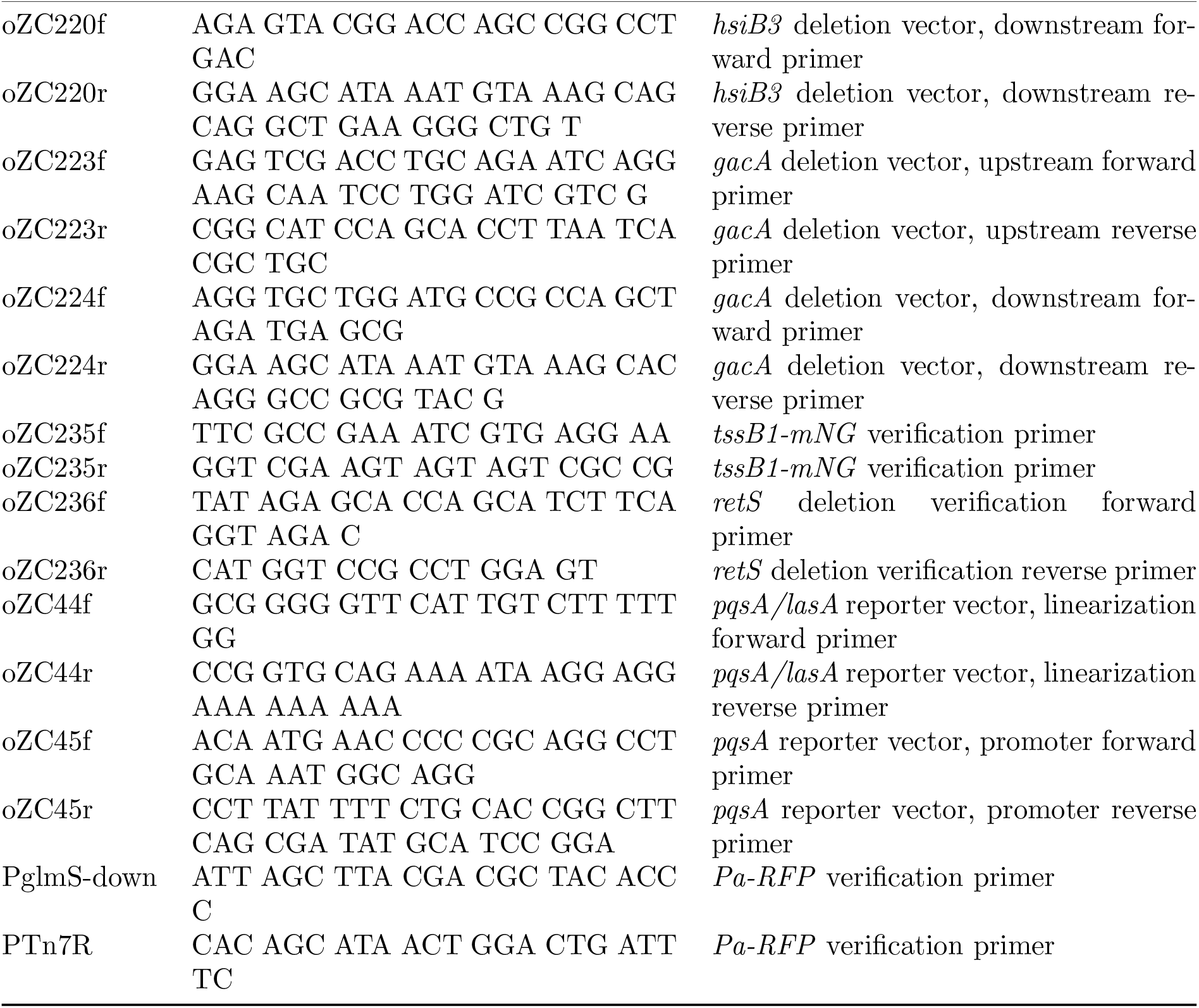
Primers used in this study.

### 5.2 Mutant and reporter strain construction

#### *P. aeruginosa* deletion strain construction

Deletion mutants of *P. aeruginosa* were generated using a modified two-step allele exchange protocol adapted from Hmelo *et al*. [48]. Primers flanking the upstream and downstream regions of each gene of interest were synthesized ([Millipore Sigma](https://www.sigmaaldrich.com/US/en)). The corresponding DNA fragments were amplified by PCR using PrimeSTAR GXL premix ([Takara, cat# R052A](https://www.takarabio.com/products/pcr/pcr-master-mixes/complex-templates/primestar-gxl-premix-fast-dye-plus)). The pEXG2 suicide vector was linearized with HindIII-HF (NEB, cat# 3104). Allele exchange constructs were assembled by Gibson assembly using Gibson Assembly Master Mix (NEB, cat# E2611) and electroporated into electrocompetent *E. coli* S17. Positive S17 transformants were selected, plasmids were isolated by miniprep (QIAGEN, cat# 27104), and sequence-verified ([Plasmidsaurus](https://plasmidsaurus.com/)). Sequence-confirmed allele exchange constructs were conjugated into *P. aeruginosa*, and deletion strains were selected and validated by whole-genome sequencing ([Plasmidsaurus](https://plasmidsaurus.com/)).

#### *P. aeruginosa* transcriptional reporter strain construction

Primers flanking the promoter regions of target genes, as well as primers for linearization of the template vector, were synthesized ([Millipore Sigma](https://www.sigmaaldrich.com/US/en)). Promoter fragments and linearized vectors were generated by PCR using PrimeSTAR GXL premix ([Takara, cat# R052A](https://www.takarabio.com/products/pcr/pcr-master-mixes/complex-templates/primestar-gxl-premix-fast-dye-plus)). Reporter constructs were assembled by Gibson assembly using Gibson Assembly Master Mix (NEB, cat# E2611) and electroporated into electrocompetent *E. coli* S17. Positive S17 transformants were selected, plasmids were isolated by miniprep (QIAGEN, cat# 27104), and sequence-verified ([Plasmidsaurus](https://plasmidsaurus.com/)). Sequence-confirmed reporter plasmids were electroporated into *P. aeruginosa*, and resulting strains were selected and validated by colony PCR and fluorescence imaging.

#### *P. aeruginosa* translational reporter strain construction

Translational reporter strains of *P. aeruginosa* were generated using the same two-step allele exchange workflow described in ***P. aeruginosa* deletion strain construction**.

#### Constitutive reporter strain construction

Constitutive fluorescent reporter strains of *P. aeruginosa* (*Pa*-RFP) and *B. cenocepacia* (*Bc*-GFP) were generated by quad-parental conjugation. The pRK2013 helper strain, the pTNS2/3 strain, donor strains, and recipient strains were grown overnight in 4 mL LB with or without appropriate selective antibiotics, then washed and resuspended in 100 µL fresh LB. Equal volumes (10 µL) of each culture were mixed and spotted onto the center of an LB agar plate, followed by incubation at 37 °C for 6 h. Cells were recovered by scraping the mating spot, resuspended in PBS, and plated onto selective media (*Pa*-RFP: *Pseudomonas* Isolation Agar (Millipore Sigma, cat# 17208) supplemented with 30 µg/mL gentamicin; *Bc*-GFP: *Burkholderia cepacia* Selective Agar ([Millipore Sigma, cat# 39573 and 39643](https://www.sigmaaldrich.com/US/en/search/burkholderia-cepacia-selective-agar?focus=products&page=1&perpage=30&sort=relevance&term=Burkholderia%20cepacia%20Selective%20Agar&type=product)) supplemented with 1 mg/mL kanamycin). Resulting strains were validated by colony PCR and fluorescence imaging.

### 5.3 Competition assay

#### Agar plate spatial competition

2 µL of each overnight culture of *P. aeruginosa, S. aureus*, and *B. cenocepacia* was inoculated adjacently onto 1.5 % LB agar plates and incubated at 37 °C for 72 h. Competition assays were performed using *n* = 6 independent biological replicates. Plates were imaged using the ChemiDoc Imaging System ([BIORAD, cat# 12003153](https://www.bio-rad.com/en-us/product/chemidoc-imaging-system?ID=OI91XQ15#fragment-2)) with the Coomassie blue illumination setting. Images (tiff format) were converted to 8-bit grayscale, contrast-inverted, and analyzed using ImageJ ([version 1.54p](https://imagej.net/ij/)). Grayscale intensity profiles were extracted along a 10-pixel-wide line crossing the interface between contacting colonies.

#### Agar plate competition for competitive index

Overnight cultures of *P. aeruginosa, S. aureus* (labeled with GFP), and *B. cenocepacia* (labeled with GFP) were diluted to OD_600_ ≈ 0.1 and mixed at a 1:1 ratio (*P. aeruginosa*:competitor). The initial ratio of *P. aeruginosa* to competitor *r*_0_ was determined by counting cells of each species using fluorescence microscopy. A 10 µL aliquot of the coculture mixture was inoculated onto 1.5 % LB agar plates and incubated at 37 °C. After incubation, colonies were scraped from the agar surface and resuspended in 50 mM Tris buffer. The final ratio of *P. aeruginosa* to competitor *r*_*t*_ was then measured. The competitive index was calculated as 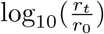 (*n* = 3).

#### Microfluidic coculture assay

Overnight cultures of fluorescently labeled *P. aeruginosa* (*Pa*-RFP), *S. aureus* (*Sa*-GFP), and *B. cenocepacia* (*Bc*-GFP) were diluted to OD_600_ ≈ 0.05, mixed at a 1:9 ratio (*Pa*:competitor), and loaded into a CellASIC ONIX plate ([Millipore Sigma, cat# B04A-03](https://www.sigmaaldrich.com/US/en/product/mm/b04a03)). The plate was controlled by CellASIC ONIX2 Microfluidic System ([Millipore Sigma, cat# CAX2-S0000](https://www.sigmaaldrich.com/US/en/product/mm/cax2s0000)), with a temperature at 37 °C and a continuous flow of fresh LB with a fluid pressure at 6.9 kPa. Time-lapse imaging was performed at 37 °C every 10 min for 24 h. GFP signal was quantified once the field of view became fully occupied and GFP intensity began to decrease. GFP area was defined as the number of pixels exceeding a threshold and normalized to the initial time point.

#### 96-well plate competition

Overnight cultures of fluorescently labeled *P. aeruginosa* (*Pa*-RFP), *S. aureus* (*Sa*-GFP), and *B. cenocepacia* (*Bc*-GFP) were diluted to OD_600_ ≈ 0.05, mixed at a 1:9 ratio (*Pa*:competitor). 200 µL of each monoculture and coculture (*n* = 9) were added into a 96-well plate (Corning, prod# 3842). The plate was incubated in an Infinite PRO 200 plate reader ([Tecan Life Sciences](https://lifesciences.tecan.com/plate_readers/infinite_200_pro?p=tab--1)) at 37 °C and 200-rpm orbital shaking. OD_600_, GFP (*λ*_*ex*_ = 480 *nm, λ*_*em*_ = 510 *nm*), and RFP (*λ*_*ex*_ = 588 *nm, λ*_*em*_ = 633 *nm*) signals were gathered every 30 min for 24 h.

### 5.4 Reporter assay

#### *P. aeruginosa* transcriptional reporter strain assay

Overnight cultures of *lasA* or *pqsA* transcriptional reporter strains were diluted in LB medium supplemented with 200 µg/mL carbenicillin (Millipore Sigma, cat# C1389) to an initial density of OD_600_ 0.05. Cells were loaded into CellASIC microfluidic plates and perfused with diluted *S. aureus* overnight culture spent medium, prepared by mixing fresh LB and spent medium at a 3:1 ratio. Plates were controlled and imaged using the same conditions described above.

#### *P. aeruginosa* translational reporter strain assay

Overnight cultures of the *P. aeruginosa* translational reporter strain and *B. cenocepacia* were diluted in LB medium to an initial density of OD_600_ ≈ 0.05. *P. aeruginosa* was then mixed with either *B. cenocepacia* or plain LB at a 1:3 ratio (*P. aeruginosa*:*B. cenocepacia* or LB), loaded into CellASIC microfluidic plates, and perfused with LB medium. Plates were controlled and imaged under the same conditions described above.

### 5.5 RNA extraction, sequencing and differential expression analysis

#### Sample preparation

Overnight cultures of *P. aeruginosa, S. aureus*, and *B. cenocepacia* were diluted and grown to exponential phase by incubating at 37 °C with 200 rpm shaking for 2 h. Monocultures were then adjusted to OD_600_ ≈ 0.05. For cocultures, *P. aeruginosa* was mixed with a competitor at a 1:4 ratio. For monoculture controls, *P. aeruginosa* was mixed with LB at the same 1:4 ratio. Cultures were subsequently incubated at 37 °C with 200 rpm shaking.

#### Total RNA extraction

Total RNA was extracted from cocultures and monocultures after 2.5 h and 20 h of incubation using the RNeasy Mini Kit ([QIAGEN, cat# 74104](https://www.qiagen.com/us/products/discovery-and-translational-research/dna-rna-purification/rna-purification/total-rna/rneasy-kits)). Briefly, 1 mL of cell culture was stabilized with RNAprotect Bacteria Reagent (QIAGEN, cat# 76506) and pelleted by centrifugation. Cell pellets were subjected to enzymatic lysis (lysozyme (Millipore Sigma, cat# SAE0152) and lysostaphin (Millipore Sigma, cat# SAE0091)) followed by proteinase K digestion. Total RNA was purified according to the manufacturer’s instructions and eluted in nuclease-free water. For each condition, *n* = 3 independent biological replicates were collected.

#### mRNA enrichment and RNA sequencing

DNA and rRNA depletion and Illumina RNA sequencing were performed by [SeqCenter](https://www.seqcenter.com/).

#### Differential expression analysis

Differential expression analysis was performed using custom scripts (code available upon request). Raw sequencing reads were quality assessed using FastQC [49] and aggregated with MultiQC [50]. Reads were aligned to the concatenated multispecies reference genome using HISAT2 [51], and gene-level read counts were generated using featureCounts [52] and HTSeq [53]. Transcript-level quantification was additionally performed using Salmon[54]. Differential expression analysis was conducted in Python using PyDESeq2 [55], with *P. aeruginosa* monoculture samples used as the reference condition. Gene set enrichment analysis was performed using GSEApy [23]. Gene annotations and pathway definitions were obtained from the NCBI database [7] and the Kyoto Encyclopedia of Genes and Genomes (KEGG [24]) database.

#### Data availability

Data are accessible at NCBI GEO database, accession GSE327547, GSE327423.

### 5.6 Genome DNA extraction and sequencing

#### Genomic DNA extraction

Genomic DNA was extracted using the DNeasy Blood and Tissue Kit ([QIA-GEN, cat# 69504](https://www.qiagen.com/us/products/discovery-and-translational-research/dna-rna-purification/dna-purification/genomic-dna/dneasy-blood-and-tissue-kit)). Briefly, 1 mL of overnight culture was pelleted and lysed according to the manufacturer’s protocol. Genomic DNA was purified and eluted in Buffer AE supplied with the kit.

#### Whole-genome sequencing and analysis

Whole-genome sequencing using Oxford Nanopore Technologies (ONT) platform and genome assembly was performed by [Plasmidsaurus](https://plasmidsaurus.com/). Analysis was performed using [Snapgene V8](https://www.snapgene.com/updates).

### 5.7 Heat-killed *S. aureus* lysis assay

#### Heat-killed *S. aureus* sample preparation

Overnight culture of *S. aureus* was washed and resuspended in 50 mM Tris buffer, and then incubated at 75 °C for 30 min.

#### Heat-killed *S. aureus* lysis assay

Serial dilutions of heat-killed *S. aureus* (HKSA) were prepared in 50 mM Tris buffer. Serial dilutions of *P. aeruginosa*:*S. aureus* coculture spent medium were generated by mixing wild-type (*P. aeruginosa*) coculture spent medium with Δ*lasA* (*P. aeruginosa*) coculture spent medium. In a 96-well plate, 80 µL of HKSA suspension was combined with 20 µL of spent medium. The plate was incubated at 37 °C with 200 rpm shaking, and OD_600_ was measured every 90 s for 6 h.

### 5.8 Simulation models

#### Spatial contact-dependent-competition simulation

Contact-dependent antagonism between *P. aeruginosa* (*Pa*) and a rival is modeled using a spatially explicit 2D agent-based framework (CellModeller [56]) with cell growth and division, mechanical interactions, stochastic contact-dependent killing, and local replacement [57].

Cells are represented as growing and dividing rods in a two-dimensional domain with a fixed timestep Δ*t* = 0.01 h. Each simulation is run for 15 000 timesteps. Each species grows with intrinsic rates *α*_*P a*_ (h^−1^) and *α*_rival_ (h^−1^), modulated by a global density-dependent slowdown reflecting the effect of crowding on competition for resources,

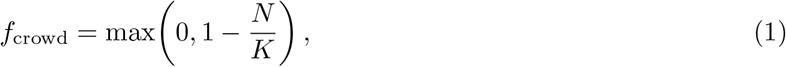

where *N* is the total population size and *K* = 40 000 is a carrying capacity. Effective growth rates are

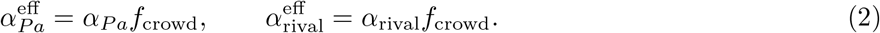

Contact-mediated killing by *Pa* is implemented as an intrinsic Poisson firing process for each *Pa* cell.

The effective growth-rate-dependent firing rate (h^−1^) is

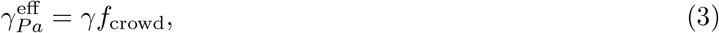

yielding a per-timestep firing probability

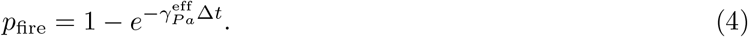

Upon firing, the *Pa* cell identifies all neighboring rival cells in direct physical contact. If at least one such neighbor is present, one is randomly selected and converted into a non-growing dead intermediate state, while retaining its original size and spatial position.

Dead cells subsequently enter a local, contact-dependent replacement process. For each dead cell, let *n*_*P a*_ and *n*_rival_ denote the numbers of neighboring *Pa* and rival cells, respectively. Species-specific replacement rates (h^−1^) are defined as

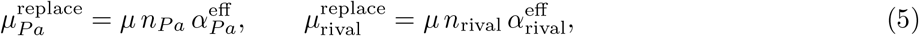

where *µ* is a dimensionless constant. Replacement occurs with probability

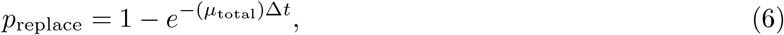

where

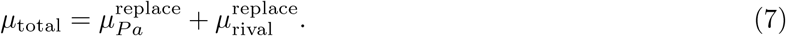

The replacement species identity is assigned proportionally to the corresponding rates,

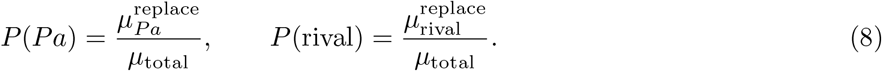

The replaced cell adopts the assigned species identity and resumes growth and division.

#### Bulk, secreted-molecule-dependent competition model

Antagonism between *P. aeruginosa* (*Pa*) and *S. aureus* (*Sa*) is modeled using a well-mixed ordinary differential equation (ODE) framework with a shared limiting resource, density-dependent slowdown of growth, and growth-coupled allocation of antagonistic-factor production by *Pa*. The state variables are *Pa* density *P*(*t*) (cell*/*L), *Sa* density *S*(*t*) (cell/L), inhibitor concentration *I*(*t*) (mol L^−1^), lytic enzyme concentration *E*(*t*) (mol L^−1^), and resource concentration *R*(*t*). Resource limitation is described by a Monod factor

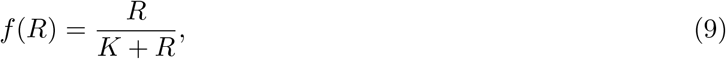

and an additional density-dependent slowdown of growth is implemented as

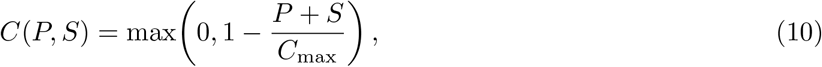

where *C*_max_ = 10^12^ cell/L is the maximum total cell density. Let *α*_*P a*_ (h^−1^) denote the intrinsic *Pa* growth rate (Supplementary Note 1). The potential *Pa* growth flux per hour is

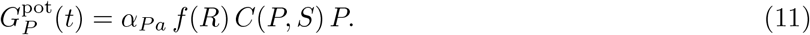

However, *Pa* dynamically allocates a fraction of its growth potential to inhibitor and lytic enzyme production through timing gates Γ_*I*_ (*t*) and Γ_*E*_(*t*) (sigmoidal activation functions). Let *t*_*X*_ denote the half-maximum onset time of inhibitor (*t*_*I*_) or lytic enzyme (*t*_*E*_) production, and *A*_*X*_ denote the allocation factor, with time course of allocation given by

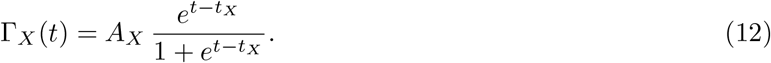

The total allocation fraction is

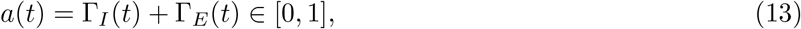

so the realized *Pa* growth flux per hour is

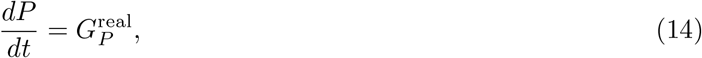

where

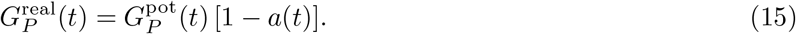

Antagonistic-factor production is assumed to be growth-coupled and proportional to *Pa* metabolic activity. In terms of the growth fluxes allocated to production, the dynamics of inhibitor *I* and lytic enzyme *E* are

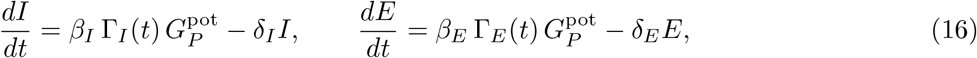

where *δ*_*I*_ (h^−1^) and *δ*_*E*_ (h^−1^) are decay rates (Supplementary Note 1). We obtain the production coefficients *β*_*I*_ and *β*_*E*_ (mol L^−1^) by converting per-division total ATP budget into maximal molar production rates based on estimated per-molecule synthesis costs (Supplementary Note 1). Inhibitor action reduces *Sa* growth through a Hill-type suppression function

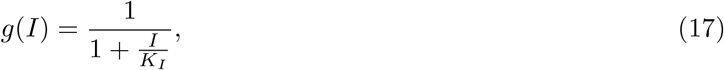

where *K*_*I*_ (mol L^−1^) is the half-inhibitory concentration (Supplementary Note 1), so that the realized *Sa* growth flux per hour is

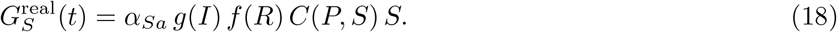

LasA-mediated killing of *Sa* is modeled using a quasi-steady-state approximation (QSSA) enzyme–substrate module. Conceptually, LasA binds an accessible wall-substrate pool associated with live *Sa*, turns over that substrate, and the resulting wall-cleavage flux is converted into an *Sa* loss term. We assume that the concentration of accessible LasA cleavage sites *B* is proportional to *S. aureus* density,

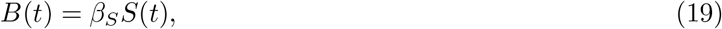

where *β*_*S*_ (mol/cell) is the effective concentration of accessible wall substrate per *S. aureus* cell (Supplementary Note 1). LasA action is modeled as

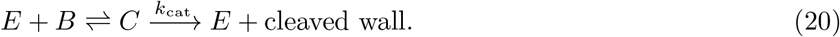

Under the exact QSSA, the transient complex concentration is then

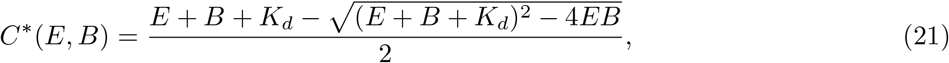

where *K*_*d*_ (mol L^−1^) is the effective dissociation constant for LasA binding to an accessible wall substrate (Supplementary Note 1). The corresponding wall-cleavage flux is

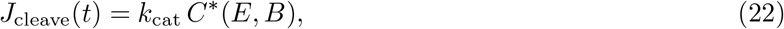

where *k*_cat_ (h^−1^) is the effective catalytic turnover rate (Supplementary Note 1). This cleavage flux is converted to *Sa* cell loss through

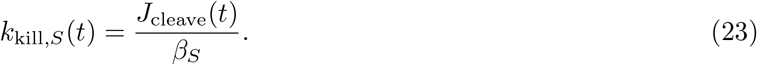

The overall *Sa* dynamics are therefore

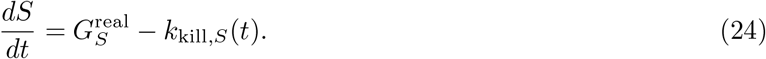

Resource depletion is directly coupled to biomass production,

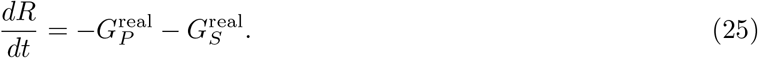

The coupled ODE system is integrated using an adaptive Runge-Kutta solver (RK45) over *t*_total_ = 24 h. Different antagonistic strategies (e.g., inhibitor-first then lytic enzyme, or lytic-enzyme-only) are implemented by varying the activation times and allocation amplitudes of Γ_*I*_ (*t*) and Γ_*E*_(*t*).

## 6 Acknowledgements

We thank Dr. Eric Déziel’s and Dr. Marek Basler’s groups for providing strains used in this research. Research reported in this publication was supported by the National Institute of General Medical Sciences (NIGMS) of the National Institutes of Health (NIH) under award numbers R01GM082938 and R01GM143227, and by the National Institute of Allergy and Infectious Diseases (NIAID) of the National Institutes of Health (NIH) under award number 5DP1AI190418. The content is solely the responsibility of the authors and does not necessarily represent the official views of the National Institutes of Health. This work was supported in part by Princeton University through the Center for the Physics of Biological Function.

## 7 Supplementary Information

### 7.1 Supplementary Tables

### 7.2 Supplementary Note 1 Parameter estimation for simulation models

#### Determination of growth rates of *Pa, Bc*, and *Sa*

Overnight cultures of *Pa, Bc*, and *Sa* were diluted 200-fold with fresh LB and incubated in a plate reader (200 µL per well in a 96-well plate) at 37 °C with 200 rpm shaking. OD_600_ was measured at 10-min intervals for 24 h. Growth rates were measured from the mid-log phase of resulting growth curves as follows (mean ± s.d., *n* = 9): *α*_*P a*_ = 0.62 ± 0.01 h^−1^, *α*_*Bc*_ = 0.60 ± 0.02 h^−1^, *α*_*Sa*_ = 0.86 ± 0.02 h^−1^.

#### Determination of decay rates *δ* of inhibitor and lytic enzyme

Decay rates *δ* of inhibitor and enzyme were derived from reported half-lives *τ*_1*/*2_ of HQNO and LasA according to 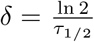. The reported *τ*_1*/*2_ values for HQNO and LasA are approximately 2 h [66] and 20 h [67], respectively.

#### Determination of production coefficients *β*_*X*_ of inhibitor and lytic enzyme

To parameterize production of the diffusible inhibitor and lytic enzyme, we conceptually convert the total cellular energy budget for biosynthesis into a maximum molecular production rate per unit biomass. The key guiding question is: if a unit density of *P. aeruginosa* were to divert all of its biosynthetic capacity into producing a single product, what would be the production rate of that product? We then scale this maximal capacity by the time-dependent allocation fractions Γ_*X*_ used in the model. Let *B*_div_ = 5 × 10^9^ATP/cell [68] denote the approximate ATP cost required to synthesize one new cell (*i*.*e*., the ATP budget per division). The division flux *G*_pot_(*t*) is defined as in Methods. Therefore the corresponding bulk ATP flux available for biosynthesis is

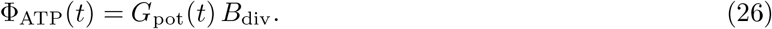

This represents the total ATP being invested into biomass production per L per hour. Let *c*_*X*_ denote the ATP cost per molecule for inhibitor or lytic enzyme (*X* ∈ {*I, E*}), if all ATP were redirected to producing *X*, the maximal molecular production flux would be 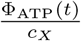. Converting to molar concentration using Avogadro’s number *N*_*A*_

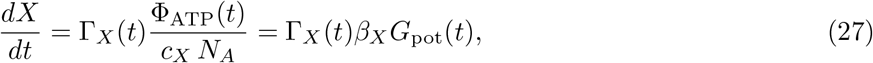

where the production coefficient is determined as

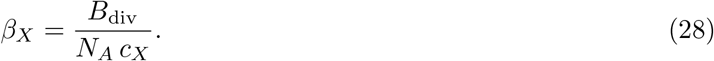

*c*_*E*_ and *c*_*I*_ were estimated using the cost of LasA and HQNO per molecule. For LasA, with *n*_aa_ amino acids, protein synthesis requires approximately 5 ATP equivalents per amino acid incorporated [69], therefore

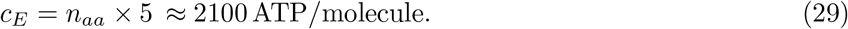

For HQNO, the dominant energetic contributions include: C7 heptyl side chain assembly via fatty-acidlike elongation (6 NAD(P)H ≈ 18 ATP), malonyl-CoA formation (~ 3 ATP), and additional redox and N-oxidation steps (~ 7 ATP equivalents) [70]. Altogether, this yields a rough estimate of

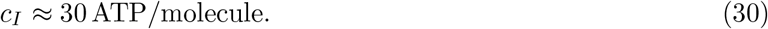

#### Determination of half-inhibitory concentration *K*_*I*_

To measure *K*_*I*_, log-phase *Sa* cultures were treated with serial diluted HQNO (*n* = 3). Growth curves for all conditions were measured using OD_600_, from which growth rates were measured. *K*_*I*_ was measured to be the HQNO concentration leading to half-maximum growth rate, which was 0.1 µg mL^−1^ or 7 × 10^−5^ mol L^−1^.

#### Determination of effective concentration of accessible wall substrate per *S. aureus* cell *β*_*S*_, effective LasA binding affinity *K*_*d*_, and catalytic turnover rate *k*_cat_

To measure *β*_*S*_, *K*_*d*_, and *k*_cat_, serial dilutions of spent medium from wild-type *P. aeruginosa* and *S. aureus* were mixed with serial dilutions of heat-killed *S. aureus* (HKSA), and lysis curves for all conditions were obtained by measuring OD_600_ at 90-s intervals for 6 h. For each condition, the matched 0% HKSA well at the same spent-medium dilution was used as background, and the background-subtracted signal was taken to represent intact HKSA material remaining over time. These lysis curves were then fit globally with the QSSA model described in the Methods section, in which LasA concentration scaled with spent-medium dilution and HKSA signal was converted to an effective substrate concentration. *β*_*S*_, *K*_*d*_, and *k*_cat_ were determined by nonlinear least-squares fitting of all time courses simultaneously across the dilution matrix. The fitted values were *β*_*S*_ = 1 × 10^−20^ mol/cell, *K*_*d*_ = 3.7 × 10^−9^ mol L^−1^, *k*_cat_ = 2.8 h^−1^.

---

**Extended Data Fig.1.**
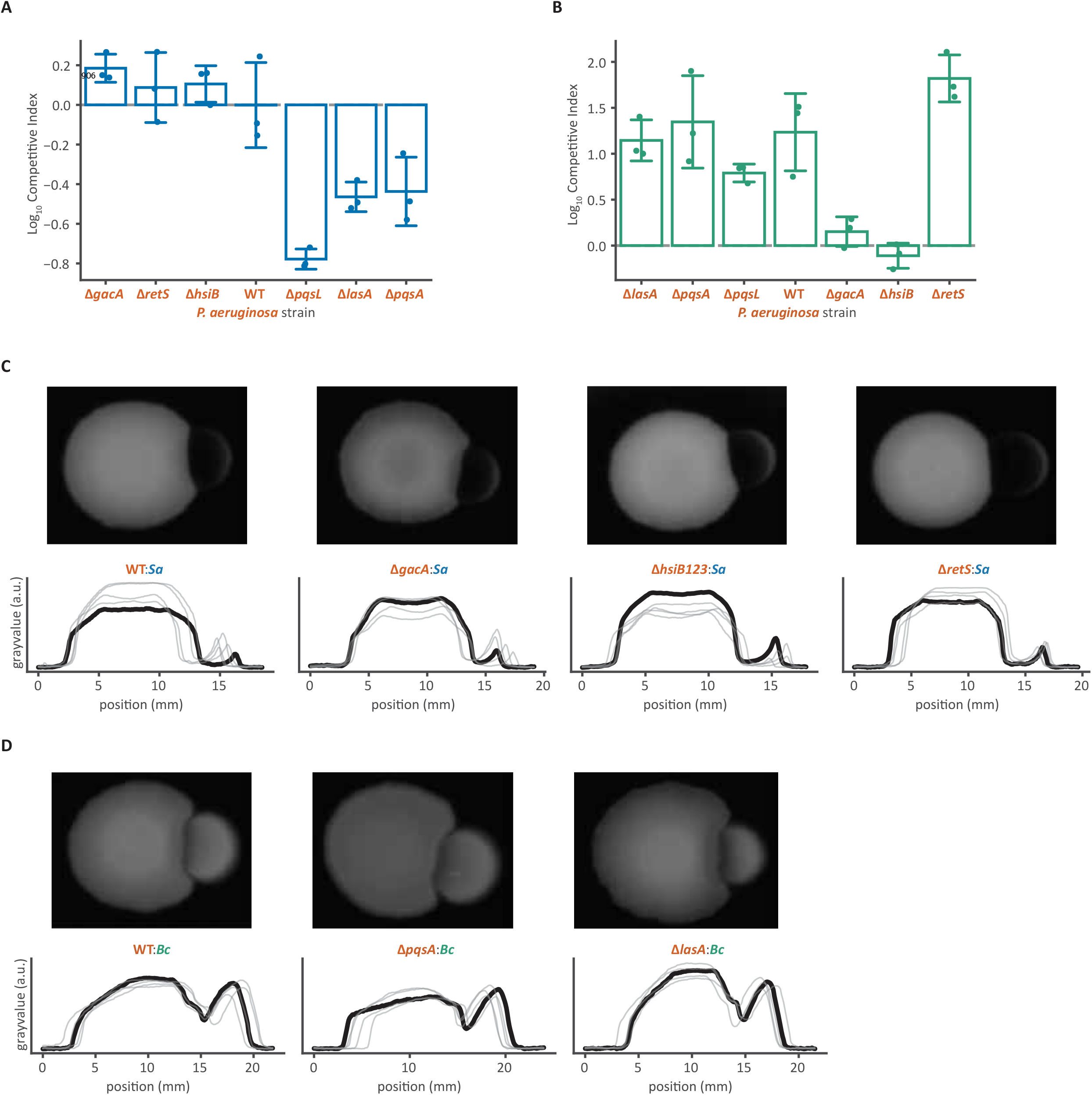
Specificity of different antagonisms. (A) Competitive indices (see Methods) of *P. aeruginosa* strains against *S. aureus*. Competitive indices were calculated after 6 h of coculture at 37 °C (*n* = 3). (B) Competitive indices of *P. aeruginosa* strains against *B. cenocepacia*. Competitive indices were calculated after 6 h of coculture at 37 °C (*n* = 3). (C) Agar plate competition assays between *P. aeruginosa* strains and *S. aureus*. Representative images (top) and corresponding grayscale intensity profiles across colonies (bottom; *n* = 5) are shown. (D) Agar plate competition assays between *P. aeruginosa* strains and *B. cenocepacia*. Representative images (top) and corresponding grayscale intensity profiles across colonies (bottom; *n* = 5) are shown.

**Extended Data Fig.2.**
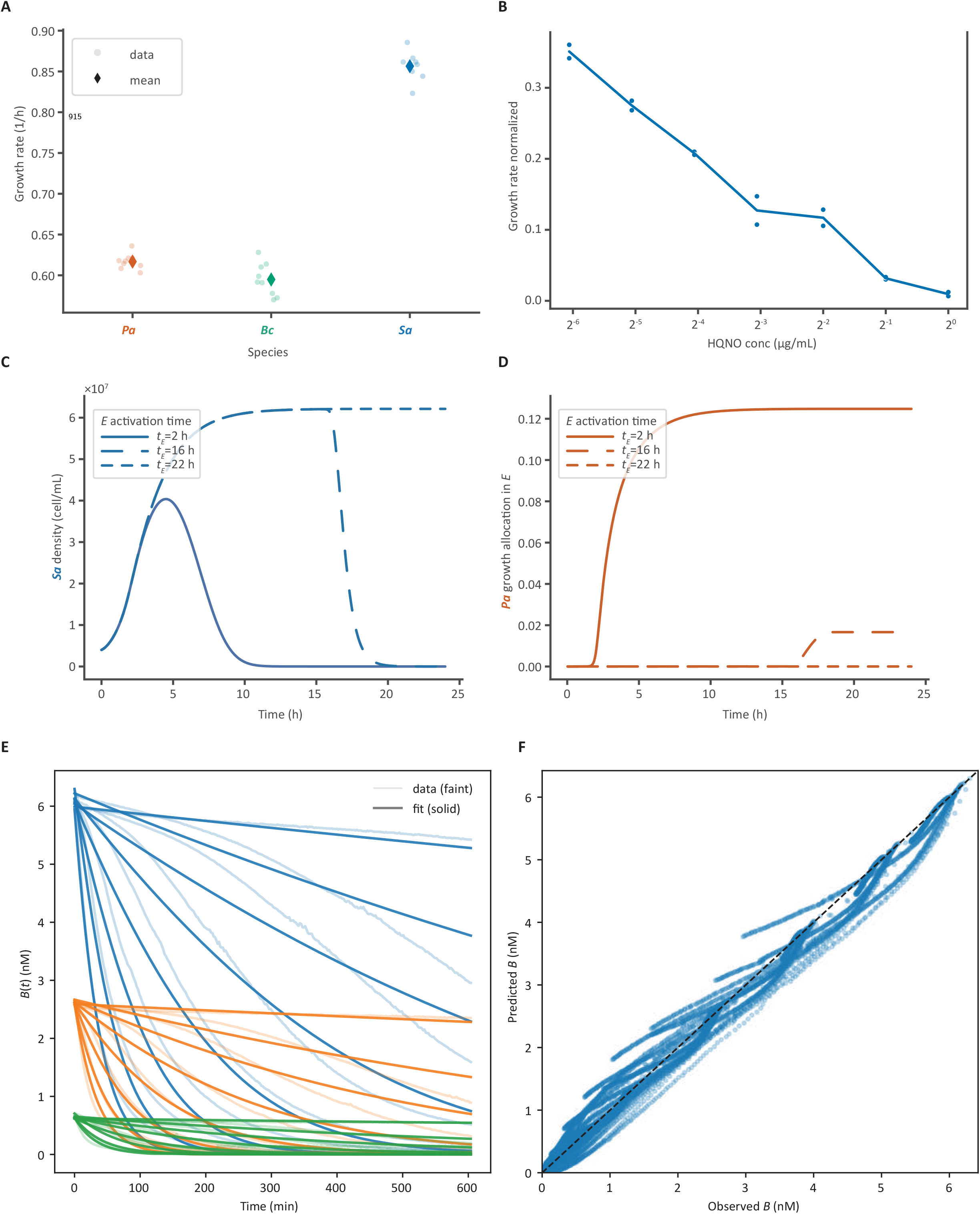
Additional analyses for the bulk, diffusible-molecule-dependent competition model. (A) Growth rates of *P. aeruginosa, B. cenocepacia*, and *S. aureus*. Monocultures of each species (*n* = 9) were initiated in a 96-well plate and incubated in a plate reader with shaking (200 rpm) at 37 °C. OD_600_ was measured every 10 min for 20 h. Growth rate was estimated by sliding a window (70 min) across the log-transformed growth curve and finding the most linear, steepest segment during the early exponential phase (0.15 ≤ OD_600_ ≤ 0.45). (B) Effect of HQNO on *S. aureus* growth. Monocultures of *S. aureus* (*n* = 2) were grown for 2 h under the conditions described in (A), after which HQNO was added at varying concentrations. Growth rates were calculated as in (A). (C) Simulated growth dynamics of *S. aureus* in the diffusible competition model under three different activation times for lytic enzyme production (*t*_*E*_). (D) Metabolic cost of lytic-enzyme production of *P. aeruginosa* in the diffusible competition model for three different activation times. *Pa* growth allocation in *E* is defined as the ratio of the resource consumption rate for enzyme synthesis (h^−1^) to that for biomass production (h^−1^). (E) Model fitting to selected experimental trajectories of effective binding site concentration (converted from OD_600_) over time. Experimental data were obtained from a heat-killed *S. aureus* (HKSA) lysis assay using serial dilutions of *Pa*:*Sa* coculture spent medium and HKSA (see Methods). These data were used to estimate parameters of the diffusive competition model. (F) Global fit of the diffusive competition model to all experimental data from the HKSA lysis assay. Each point represents the predicted (model) versus observed (experimental) effective binding site concentration at a given time point under a specific condition.

**Extended Data Fig.3.**
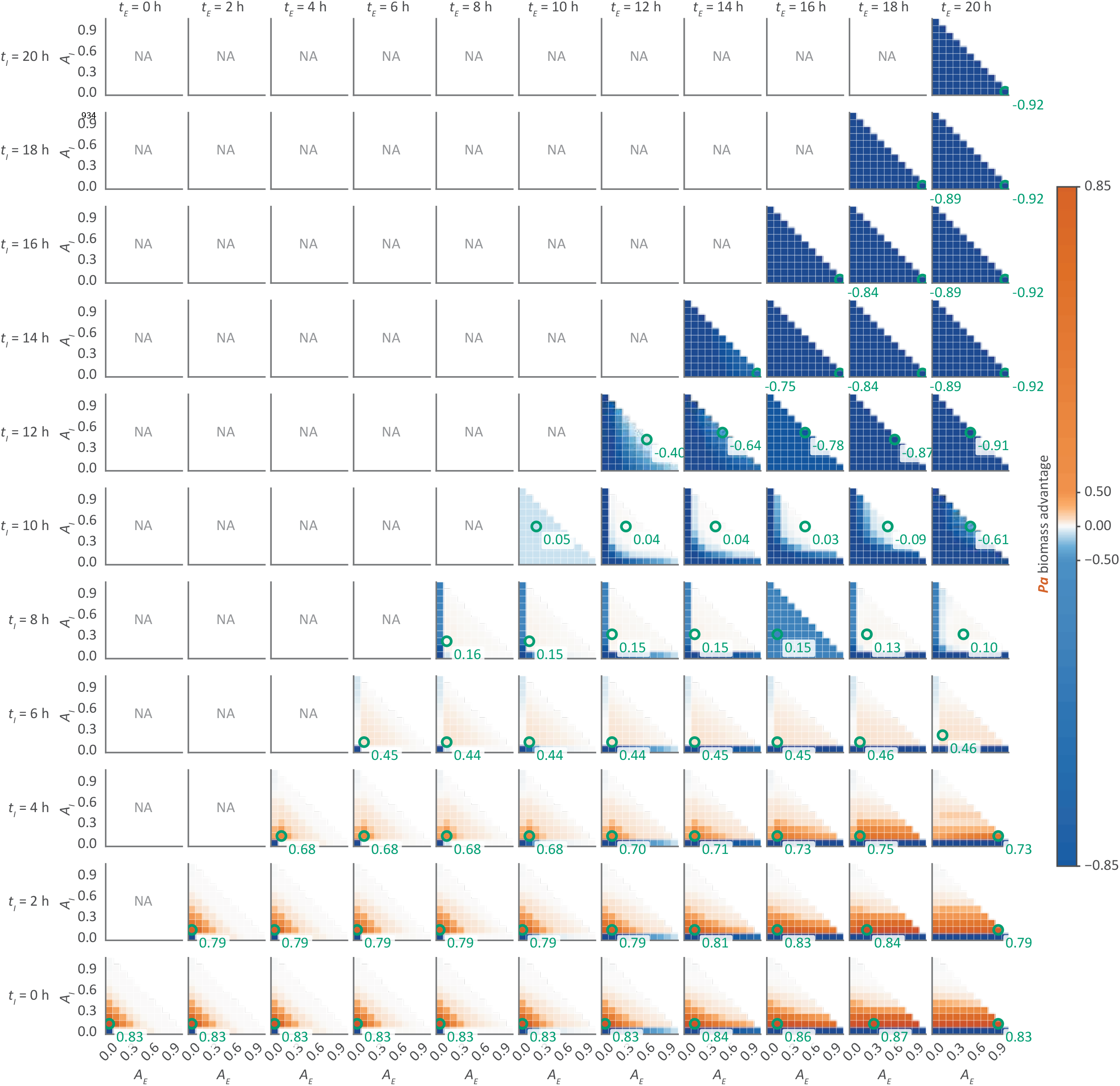
Optimization of activation times and allocation amplitudes of inhibitor and lytic-enzyme production. Each data point represents the *Pa* biomass advantage in a simulation with specific activation times (*t*_*X*_) and allocation amplitudes (*A*_*X*_) of inhibitor and lytic-enzyme production. Each heatmap has fixed *t*_*X*_ and varied *A*_*X*_, with the *A*_*I*_ and *A*_*E*_ pair leading to the maximum *Pa* biomass advantage highlighted.

**Extended Data Fig.4.**
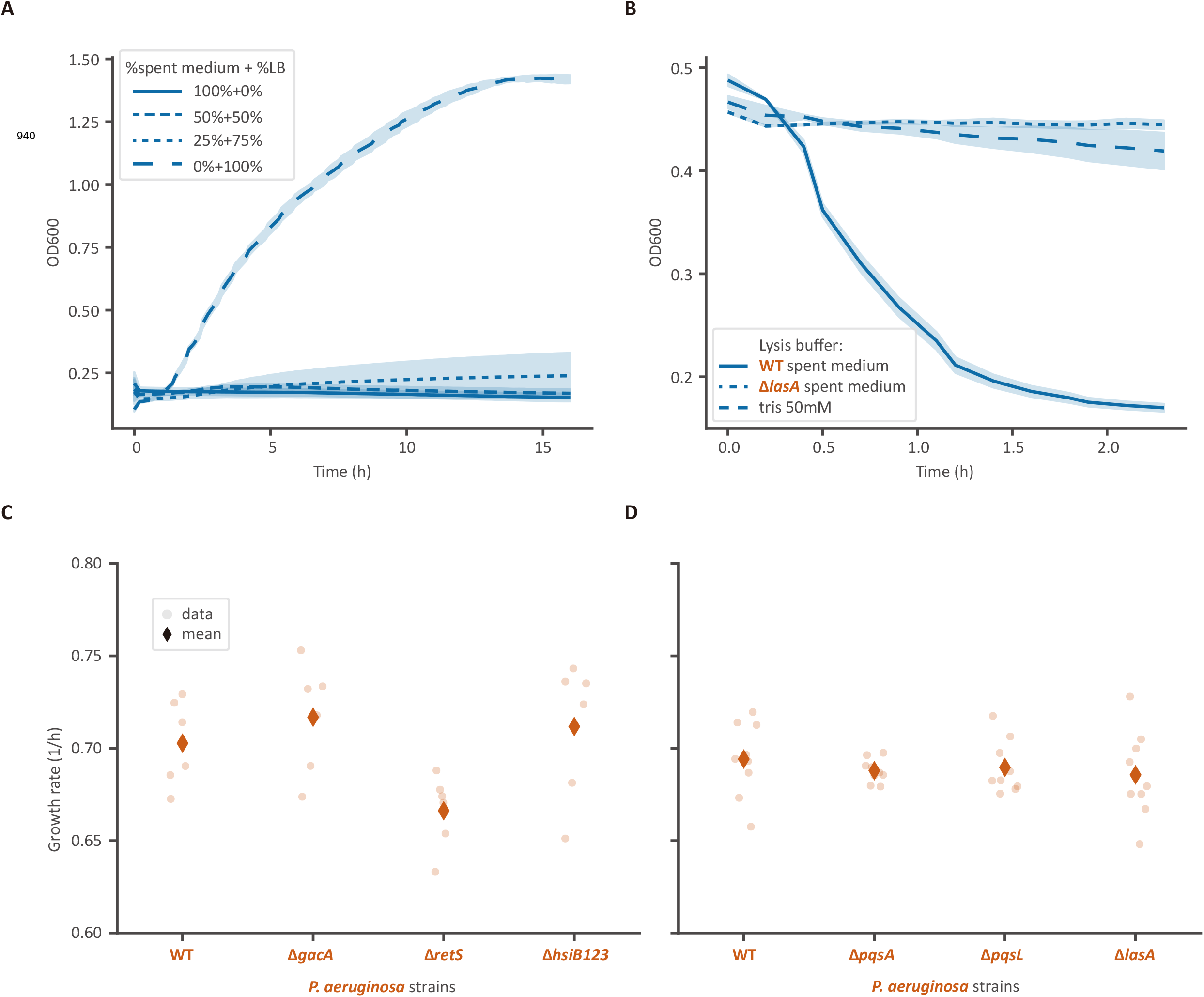
Additional results on heat-killed *S. aureus* lysis assay and *P. aeruginosa* mutant strain growth rates. (A) Growth of *S. aureus* monoculture in *Pa*:*Sa* spent medium diluted with fresh LB. Monocultures (*n* = 3) were initiated in a 96-well plate and incubated in a plate reader with shaking (200 rpm) at 37 °C. OD_600_ was measured every 10 min for 20 h. (B) Heat-killed *S. aureus* lysis assay. Spent media from different *Pa*:*Sa* cocultures (*n* = 3) were added to heat-killed *S. aureus* and OD_600_ was measured as described in (A). (C and D) Growth rates of different *P. aeruginosa* strains in monocultures. Growth curves of each strain (C: *n* = 6, D: *n* = 9) were measured as described in (A). Growth rate was estimated by sliding a window (70 min) across the log-transformed growth curve and finding the most linear, steepest segment during the early exponential phase (0.15 ≤ OD_600_ ≤ 0.45).

**Extended Data Movie 1 Competition of *P. aeruginosa* and *S. aureus* in microfluidic chamber**. Fluorescently labeled *P. aeruginosa* (DsRed2) and *S. aureus* (sGFP) were diluted to OD_600_ ≈ 0.05, mixed at a 1:9 ratio (*Pa*:*Sa*), and loaded into a CellASIC ONIX plate. The plate was maintained at 37 °C with a continuous flow of fresh LB. Time-lapse imaging was performed every 10 min for 24 h. In the video *P. aeruginosa* and *S. aureus* were colored as magenta and green, respectively.

**Extended Data Movie 2 Competition of *P. aeruginosa* and *B. cenocepacia* in microfluidic chamber**. Fluorescently labeled *P. aeruginosa* (DsRed2) and *B. cenocepacia* (eGFP) were diluted to OD_600_ ≈ 0.05, mixed at a 1:9 ratio (*Pa*:*Bc*), and loaded into a CellASIC ONIX plate. The plate was maintained at 37 °C with a continuous flow of fresh LB. Time-lapse imaging was performed every 10 min for 24 h. In the video *P. aeruginosa* and *B. cenocepacia* were colored as magenta and green, respectively.

